# Structural characterization of ligand binding and pH-specific enzymatic activity of mouse Acidic Mammalian Chitinase

**DOI:** 10.1101/2023.06.03.542675

**Authors:** Roberto Efraín Díaz, Andrew K. Ecker, Galen J. Correy, Pooja Asthana, Iris D. Young, Bryan Faust, Michael C. Thompson, Ian B. Seiple, Steven J. Van Dyken, Richard M. Locksley, James S. Fraser

**Affiliations:** Department of Bioengineering and Therapeutic Sciences, University of California, San Francisco, San Francisco, CA 94158, USA; Tetrad Graduate Program, University of California, San Francisco, San Francisco, CA 94158, USA; Department of Pharmaceutical Chemistry, University of California, San Francisco, San Francisco, CA 94158, USA; Cardiovascular Research Institute, University of California, San Francisco, San Francisco, CA 94158, USA; Chemistry and Chemical Biology Graduate Program, University of California, San Francisco, San Francisco, CA 94158, USA; Department of Biochemistry and Biophysics, University of California, San Francisco, San Francisco, CA 94158, USA; Biophysics Graduate Program, University of California, San Francisco, San Francisco, CA 94158, USA; Department of Chemistry and Chemical Biology, University of California, Merced, Merced, CA 95343, USA; Department of Pathology and Immunology, Washington University School of Medicine in St. Louis, St. Louis, MO 63110, USA; Department of Medicine, University of California, San Francisco, San Francisco, CA 94158, USA; Department of Microbiology and Immunology, University of California, San Francisco, San Francisco, CA 94143, USA; University of California, San Francisco, Howard Hughes Medical Institute, San Francisco, CA 94143, USA

## Abstract

Chitin is an abundant biopolymer and pathogen-associated molecular pattern that stimulates a host innate immune response. Mammals express chitin-binding and chitin-degrading proteins to remove chitin from the body. One of these proteins, Acidic Mammalian Chitinase (AMCase), is an enzyme known for its ability to function under acidic conditions in the stomach but is also active in tissues with more neutral pHs, such as the lung. Here, we used a combination of biochemical, structural, and computational modeling approaches to examine how the mouse homolog (mAMCase) can act in both acidic and neutral environments. We measured kinetic properties of mAMCase activity across a broad pH range, quantifying its unusual dual activity optima at pH 2 and 7. We also solved high resolution crystal structures of mAMCase in complex with oligomeric GlcNAcn, the building block of chitin, where we identified extensive conformational ligand heterogeneity. Leveraging these data, we conducted molecular dynamics simulations that suggest how a key catalytic residue could be protonated via distinct mechanisms in each of the two environmental pH ranges. These results integrate structural, biochemical, and computational approaches to deliver a more complete understanding of the catalytic mechanism governing mAMCase activity at different pH. Engineering proteins with tunable pH optima may provide new opportunities to develop improved enzyme variants, including AMCase, for therapeutic purposes in chitin degradation.

## Introduction

Chitin, a polymer of β(1-4)-linked *N*-acetyl-D-glucosamine (GlcNAc), is the second most abundant polysaccharide in nature. Chitin is present in numerous pathogens, such as nematode parasites, dust mites, and fungi^1–3^, and is a pathogen-associated molecular pattern (PAMP) that activates mammalian innate immunity^4^. To mitigate constant exposure to environmental chitin, mammals have evolved unusual multi-gene loci that are highly conserved and encode chitin-response machinery, including chitin-binding (chi-lectins) and chitin-degrading (chitinases) proteins.

Humans express two active chitinases as well as five chitin-binding proteins that recognize chitin across many tissues^7^. Chitin levels can be potentially important for mammalian lung and gastrointestinal health. These tissues have distinct pH, with the lung environment normally ∼pH 7.0 and the stomach environment normally ∼pH 2.0, which raises the question of how chitin-response machinery has evolved to function optimally across such diverse chemical environments. Acidic Mammalian Chitinase (AMCase, also known as Chia, for chitinase, acidic) was originally discovered in the stomach and named for its acidic isoelectric point. AMCase is also constitutively expressed in the lungs at low levels and overexpressed upon chitin exposure^6,8,9^, suggesting this single enzyme has evolved to perform its function under vastly different chemical conditions. Chitin clearance is particularly important for mammalian pulmonary health, where exposure to and accumulation of chitin can be deleterious. In the absence of AMCase, chitin accumulates in the airways, leading to epithelial stress, chronic activation of type 2 immunity, and age-related pulmonary fibrosis^5,6^.

AMCase is a member of the glycosyl hydrolase family 18 (GH18)^10^, the members of which hydrolyze sugar linkages through a conserved two-step mechanism where the glycosidic oxygen is protonated by an acidic residue and a nucleophile adds into the anomeric carbon leading to elimination of the hydrolyzed product (**Figure 1**A). This mechanism is corroborated by structures of different GH18 chitinases, most notably *S. marcescens* Chitinase A (PDB ID: 1FFQ)^11^. In inhibitor-bound structures for human AMCase (hAMCase; PDB ID: 3FY1), interactions mimicking the retentive, post-cleavage intermediate state pre-hydrolysis of the oxazolinium intermediate are adopted by the nonhydrolyzable analogs^12,13^. Unlike the nonhydrolyzable inhibitors, we expect that the oxazolinium intermediate formed from chitin will reopen into the reducing-end GlcNAc monomer unit upon the nucleophilic addition of water.

**Figure 1.**
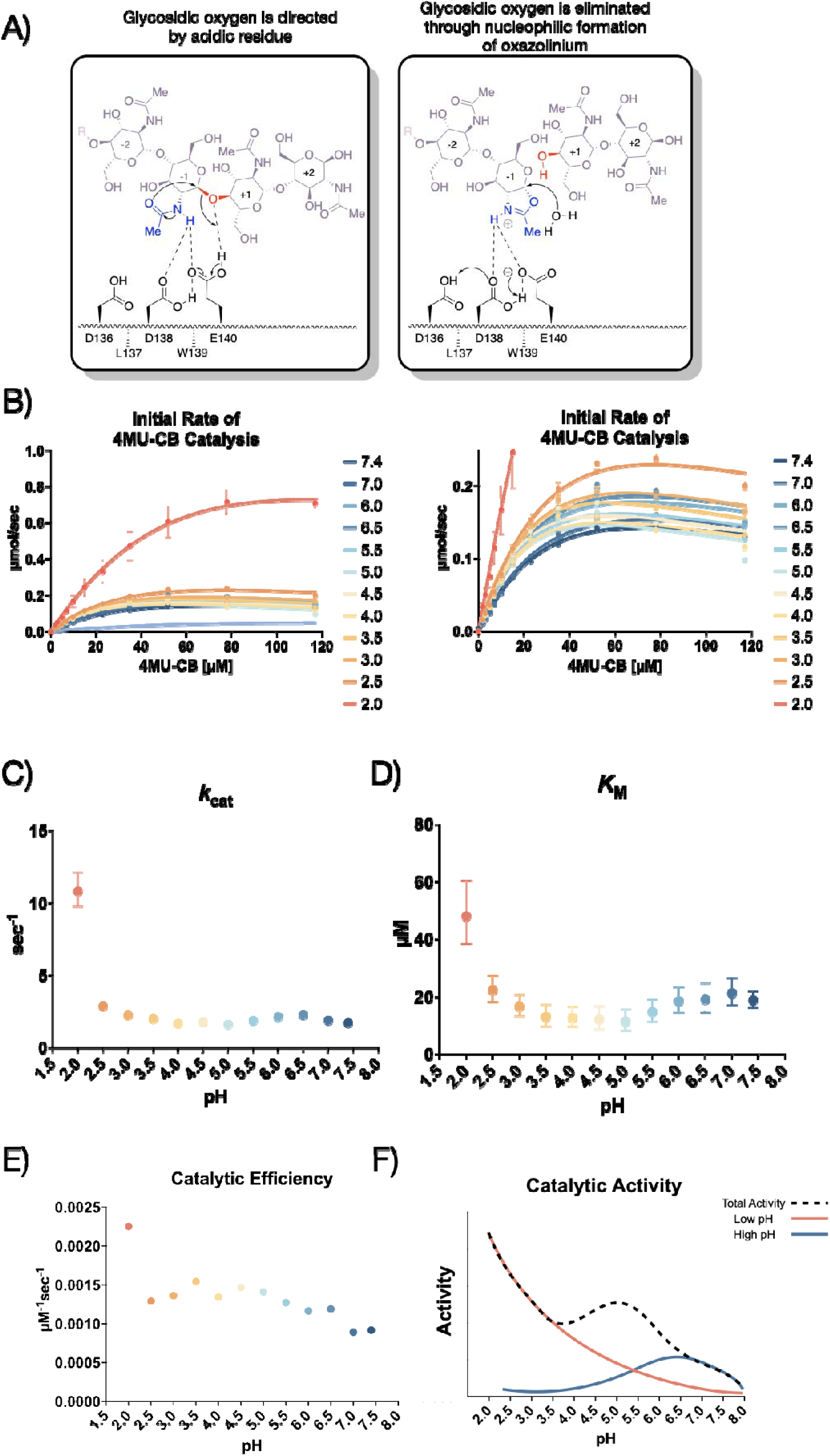
Kinetic properties of mAMCase catalytic domain at various pH. **A)** Chemical depiction of the conserved two-step mechanism where the glycosidic oxygen is protonated by an acidic residue and a nucleophile adds into the anomeric carbon leading to elimination of the hydrolyzed product. **B)** The rate of 4MU-chitobioside catalysis (1/sec) by mAMCase catalytic domain is plotted as a function of 4MU-chitobioside concentration (µM). Each data point represents n = 4 with error bars representing the standard deviation. Michaelis-Menten equation without substrate inhibition was used to estimate the *k*_cat_ and *K*M from the initial rate of reaction at various substrate concentrations. **C)** The rate of substrate turnover (1/sec) by mAMCase catalytic domain is plotted as a function of pH. Error bars represent the 95% confidence interval. **D)** The Michaelis-Menten constant of mAMCase catalytic domain is plotted as a function of pH. Error bars represent the 95% confidence interval. **E)** The catalytic efficiency (*k*_cat_/*K*_M_) of mAMCase catalytic domain is plotted as a function of pH. **F)** Hypothetical catalytic activity modeled explained by a low pH mechanism (red), and high pH mechanism (blue) and their corresponding total activity (dashed line).

Biochemical studies of mouse AMCase (mAMCase) measuring relative activity levels demonstrated a global maximum activity at acidic pH, but also a broad second local optimum near neutral pH^14^. This result suggested that mAMCase exhibits two distinct pH optima, which is unlike most enzymes that exhibit a shift or broadening of enzymatic activity across conditions^15–17^. For mAMCase the global maximum near pH 2.0 resembles the chemical environments of the stomach and the local maximum near pH 7.0 is similar to the environment of the lung. These two pH optima in the same enzyme suggest that mAMCase may employ different mechanisms to perform its function in different environments^18^. In contrast, the human homolog has maximal activity at pH 4.6 with sharply declining activity at more acidic and basic pH^18,19^. This optimum corresponds with the pH of lung tissue in pulmonary fibrosis and other disease contexts, suggesting that hAMCase may have been selected for its ability to clear chitin from the lungs and restore healthy lung function.

The activity of mAMCase has been previously measured through endpoint experiments with limited insight into the rate of catalysis, substrate affinity, and potential substrate inhibition^18^. While the pH profile of mAMCase has been reported as a percentage of maximum activity at a given pH, it is unclear how the individual kinetic parameters (*K*M or *k*_cat_ ) vary^14^.

These gaps have made it challenging to define the mechanism by which mAMCase shows distinct enzymatic optima at different pHs. One possibility is that mAMCase undergoes structural rearrangements to support this adaptation. Alternatively AMCase may have subtly different mechanisms for protonating the catalytic glutamic acid depending on the environmental pH.

In this work, we explore these hypotheses by employing biophysical, biochemical, and computational approaches to observe and quantify mAMCase function at different pHs. We measured the mAMCase hydrolysis of chitin, which revealed significant activity increase under more acidic conditions compared to neutral or basic conditions. To understand the relationship between catalytic residue protonation state and pH-dependent enzyme activity, we calculated the theoretical pKa of the active site residues and performed molecular dynamics (MD) simulations of mAMCase at various pHs. We also directly observed conformational and chemical features of mAMCase between pH 4.74 to 5.60 by solving X-ray crystal structures of mAMCase in complex with oligomeric GlcNAcn across this range. Together these data support a model in which mAMCase employs two different mechanisms for obtaining a proton in a pH-dependent manner, providing a refined explanation as to how this enzyme recognizes its substrate in disparate environments.

## Results

### New assay confirms broad pH profile for mAMCase

Prior studies have focused on relative mAMCase activity at different pH^14,18,20^ , limiting the ability to define its enzymological properties precisely and quantitatively across conditions of interest. To expand upon these previous observations of dual optima in mAMCase activity at pH 2.0 and 7.0, we measured mAMCase activity *in vitro*. We developed an approach that would enable direct measurement of *k*cat and *K*M for mAMCase across a broad pH range by modifying a prior assay that continuously measures mAMCase-dependent breakdown of a fluorogenic chitin analog, 4-methylumbelliferone (4MU) conjugated chitobioside. To overcome the pH-dependent fluorescent properties of 4MU-chitobioside, we reverted the assay into an endpoint assay, which allowed us to measure substrate breakdown across different pH^21^ (**Supplemental Figure 1**A).

We conducted our endpoint assay across a pH range of 2.0 to 7.4 to reflect the range of physiological conditions at its *in vivo* sites of action (**Figure 1**B; Data available at doi: 10.5281/zenodo.8250616). We then derived the Michaelis-Menten parameters at each pH value measured (**Supplemental Figure 2**A-C; Data available at doi: 10.5281/zenodo.8250616). We found that mAMCase has maximum activity at pH 2.0 with a secondary local maximum at pH 6.5, pointing to a bimodal distribution of activity across pH. This is consistent with the relative activity measurements previously performed on mAMCase, but distinct from a single broad pH range, as has been observed for *k*cat of hAMCase^14,18^. The two maxima at pH 2.0 and 6.5 are an approximate match the pH at the primary *in vivo* sites of mAMCase expression, the stomach and lungs, respectively^18^. These observations raise the possibility that mAMCase, unlike other AMCase homologs, may have evolved an unusual mechanism to accommodate multiple physiological conditions.

We also found that low pH primarily improves the rate of mAMCase catalysis 6.3-fold (*k*cat; **Figure 1**C), whereas *K*M (**Figure 1**D) worsens 2.5-fold from pH 7.4 to pH 2.0. Similar to chitotriosidase the other active chitinase in mammals and also a GH18 chitinase, we observe an apparent reduction in the rate of mAMCase catalysis across all pH values measured at 4MU-chitobioside concentrations above 80 μM, which suggests that mAMCase may be subject to product inhibition^22^. The underlying mechanism for the observed product inhibition may be that mAMCase can transglycosylate the products, as has been previously observed at pH 2.0 and 7.0^23^. This potential product inhibition leads to a systematic underprediction of rates by the Michaelis-Menten model at high substrate concentrations. The catalytic efficiency (*kcat*/*K*M) of mAMCase may not capture the effects of product inhibition given that these constants reflect sub-saturating substrate concentrations. Independent of the potential for product inhibition, the trend that mAMCase has highest kcat at very low pH and another local optimum at more neutral pH is clear. We hypothesize that these activity data resemble two overlapping activity distributions, suggesting that the rate at lower pH activity is dependent on the concentration of free protons in solution and that the higher pH optimum results from a distinct mechanism (**Figure 1**E).

### Characterization of mAMCase ligand occupancy and conformational heterogeneity

Our biochemical analyses led us to hypothesize that the pH-dependent activity profile of mAMCase is linked to the mechanism by which catalytic residues are protonated. Previous structural studies on AMCase have focused on interactions between inhibitors like methylallosamidin and the catalytic domain of the protein. We built on these efforts by solving the structure of mAMCase in complex with oligomeric GlcNAcn, the building block of chitin. We used chitin oligomers because they are chemically identical to polymeric chitin found in nature but are soluble and therefore more amenable for co-crystallization than crystalline chitin is. We successfully determined high resolution X-ray crystal structures of the apo mAMCase catalytic domain at pH 5.0 and 8.0 (PDB ID: 8FG5, 8FG7) and holo mAMCase catalytic domain between pH 4.74 to 5.60 in complex with either GlcNAc_2_ or GlcNAc3 (PDB ID: 8GCA, 8FRC, 8FR9, 8FRB, 8FRD, 8FRG, 8FRA; **Supplemental Figure 3**A,B; **Table 1**).

**Table 1.**
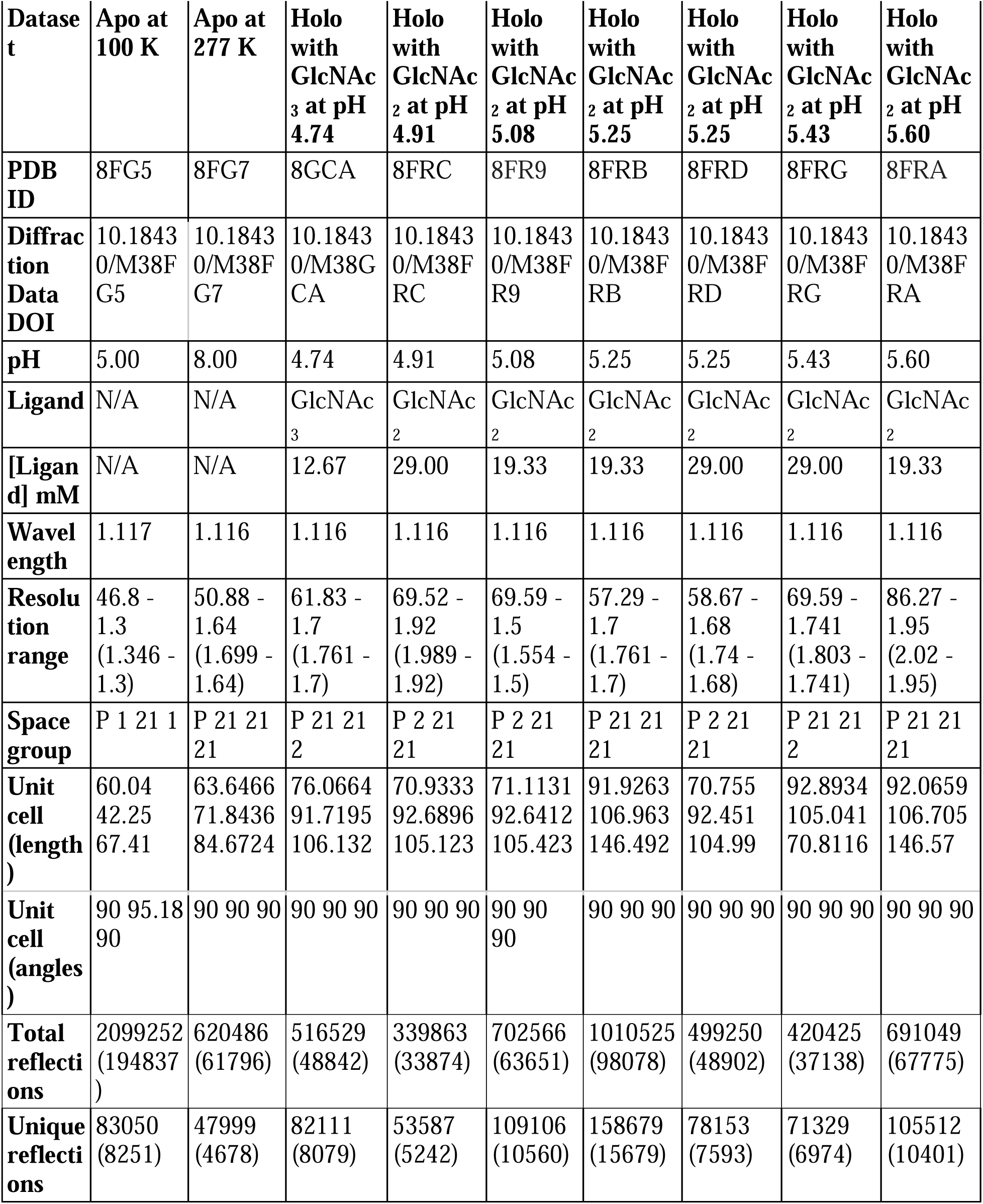
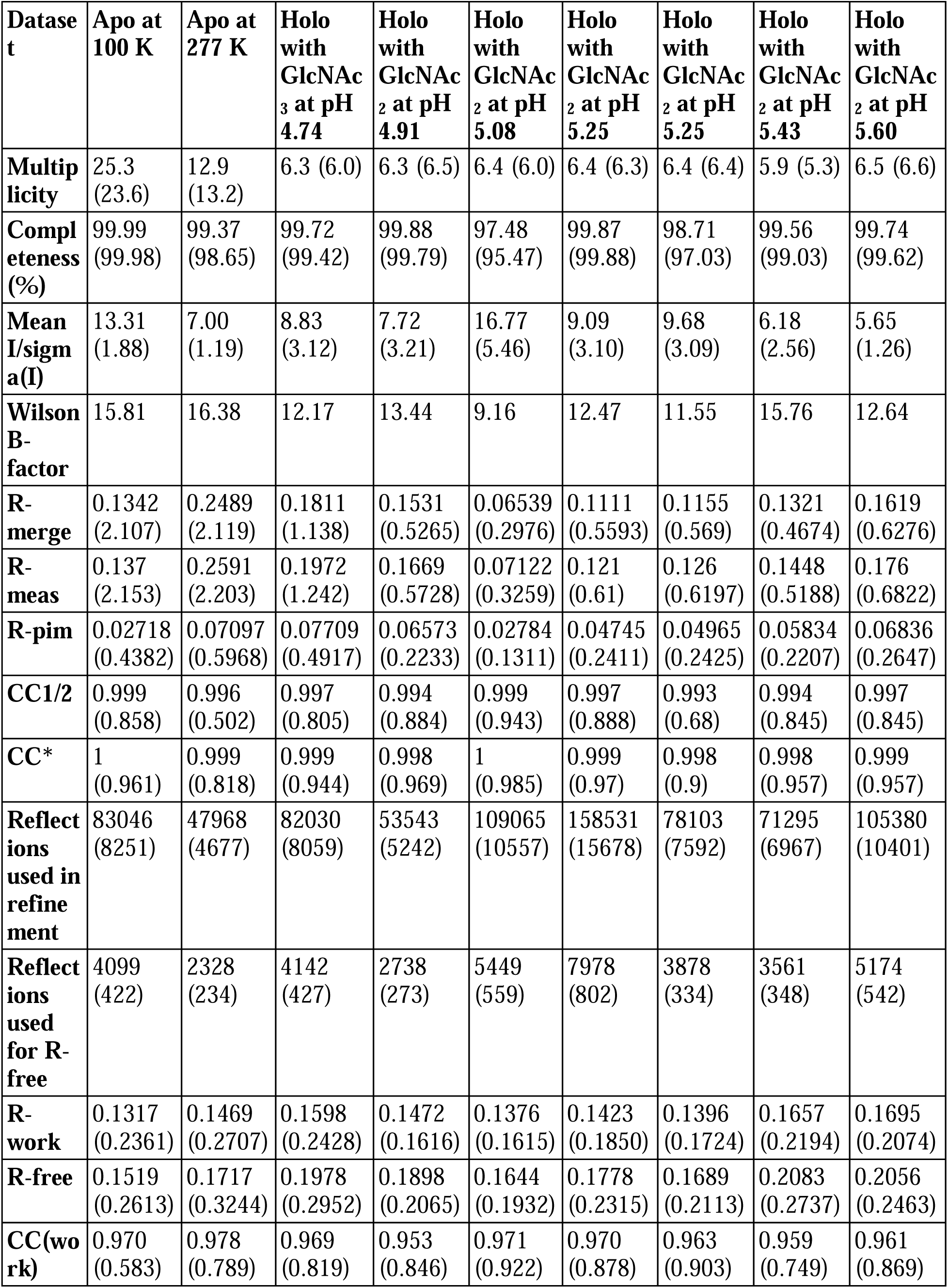
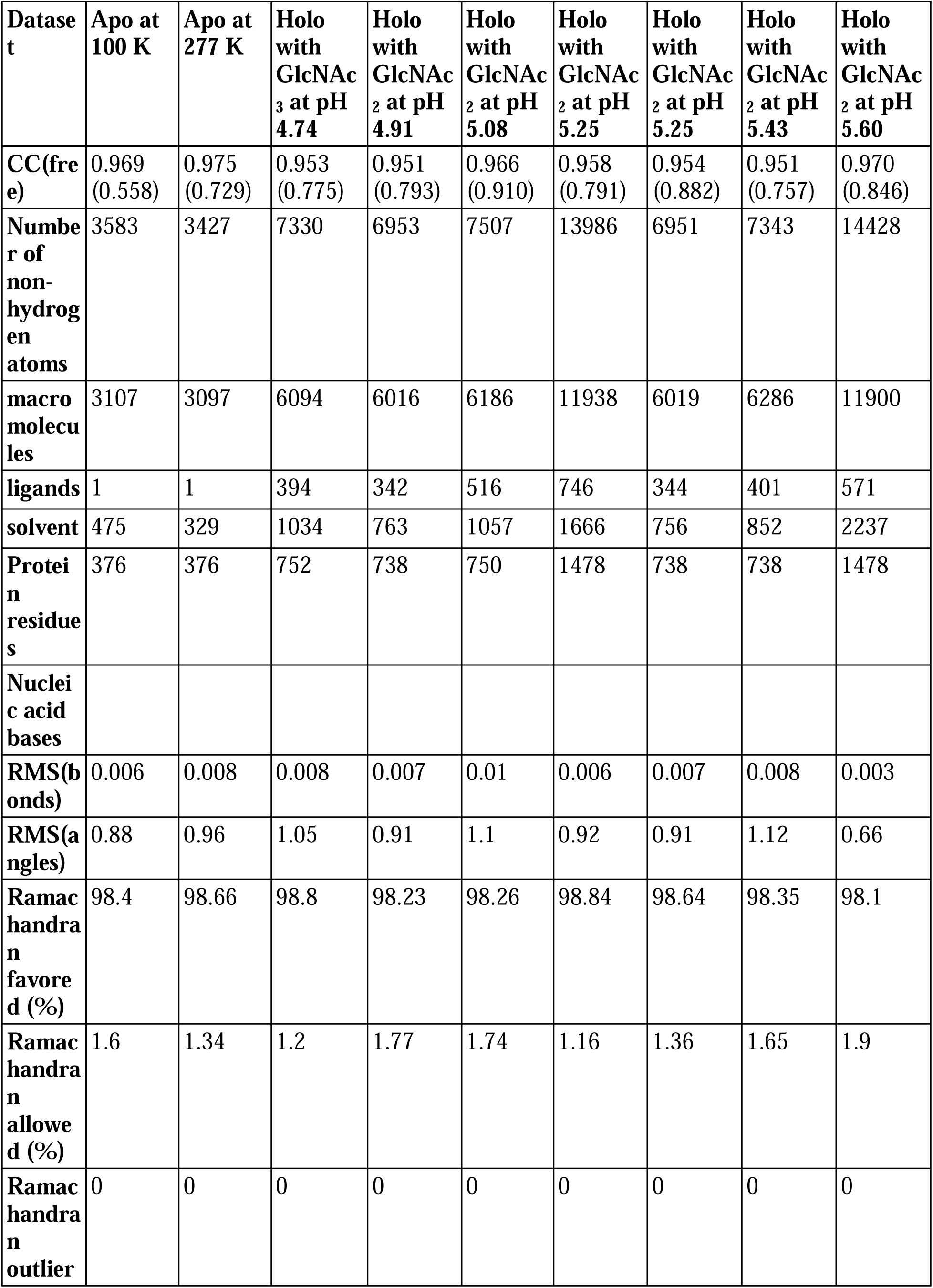
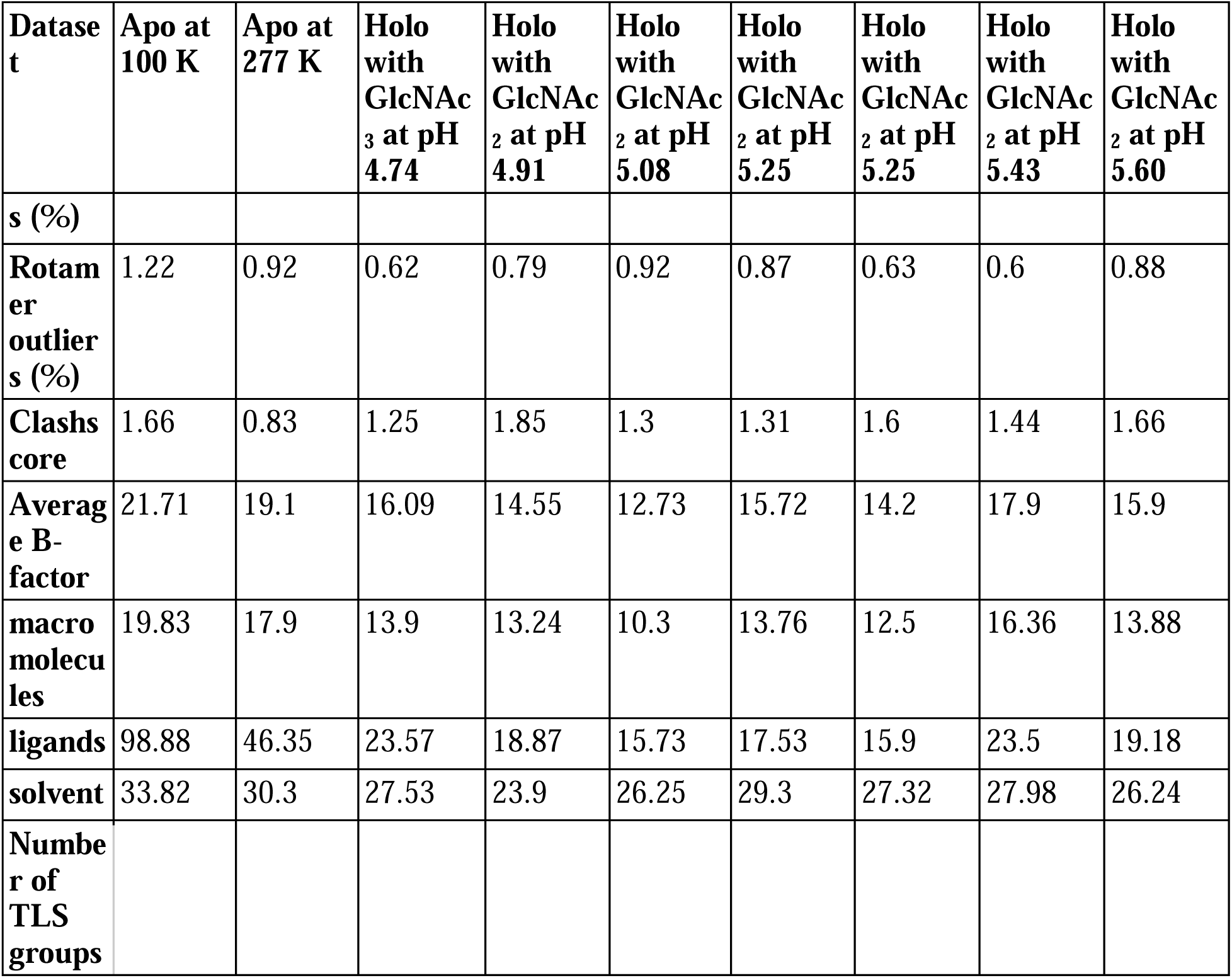
Data collection and refinement statistics. Statistics for the highest-resolution shell are shown in parentheses.

Across these different datasets we observed complex ligand density in the active site of mAMCase. In all of our datasets, we observed continuous ligand density that resembled higher order chitin oligomers (e.g. GlcNAc_4_, GlcNAc_5_, or GlcNAc_6_). This observation was confusing given that these structures were co-crystallized with either GlcNAc_2_ or GlcNAc_3_. For example, due to the continuous nature of ligand density observed in our mAMCase-GlcNAc_3_ co-crystal structure at pH 4.74 (PDB ID: 8GCA, chain A), we initially modeled hexaacetyl-chitohexaose (H-(GlcNAc)6-OH) into the -4 to +2 sugar-binding subsites, using the nomenclature for sugar-binding subsites from Davies et al.^24^. This nomenclature defines the sugar-binding subsites as *-n* to *+n*, with *-n* corresponding to the non-reducing end and *+n* the reducing end.

We next continued with a modeling approach that replaced higher order oligomer models with models that only used the chemically defined oligomers present in the crystallization drop. To accomplish this modeling of different binding poses, we placed multiple copies of these oligomers consistent with an interpretation of extensive conformational heterogeneity (**Supplemental Figure 5**D). In one sample co-crystallized with GlcNAc_3_ at pH 4.74 (PDB ID: 8GCA, chains A-B), we identified ligand density that was consistent with GlcNAc_2_, suggesting that some hydrolysis occurs in the crystal. The resulting model includes compositional heterogeneity as there are both types of oligomer present.

Therefore, across all of our datasets, we modeled a combination of ligand binding events consisting of overlapping GlcNAc_2_ or GlcNAc_3_ molecules at each sugar-binding site, i.e. GlcNAc_2_ ResID 401 Conf. A occupied subsites -3 to -2 while GlcNAc_2_ ResID 401 Conf. C occupied subsites -2 to -1. By providing each ligand molecule with an alternative conformation ID, this allowed both occupancies and B-factors to be refined (**Figure 2**A,B,C; additional details in Methods). Across these different datasets, we observed ligand density for different combinations of occupancy over the -4 to +2 sugar-binding subsites (**Figure 2**A). While modeling chito-oligomers into strong electron density, we observed strong positive difference density between sugar-binding subsites near the C2 *N*-acetyl and the C6’ alcohol moieties. Using the non-crystallographic symmetry (NCS) “ghost” feature in *Coot*, we were then able to observe that the positive difference density between ligand subsites in one chain could be explained by the dominant ligand pose observed in another associated crystallographic chain, suggesting the presence of a low-occupancy binding events. This observation led to the discovery that GlcNAcn occupies intermediate subsites, which we label *n*+0.5, continuing to follow the nomenclature established by Davies et al., in addition to canonical sugar-binding subsites (**Figure 2**B)^24^.

**Figure 2.**
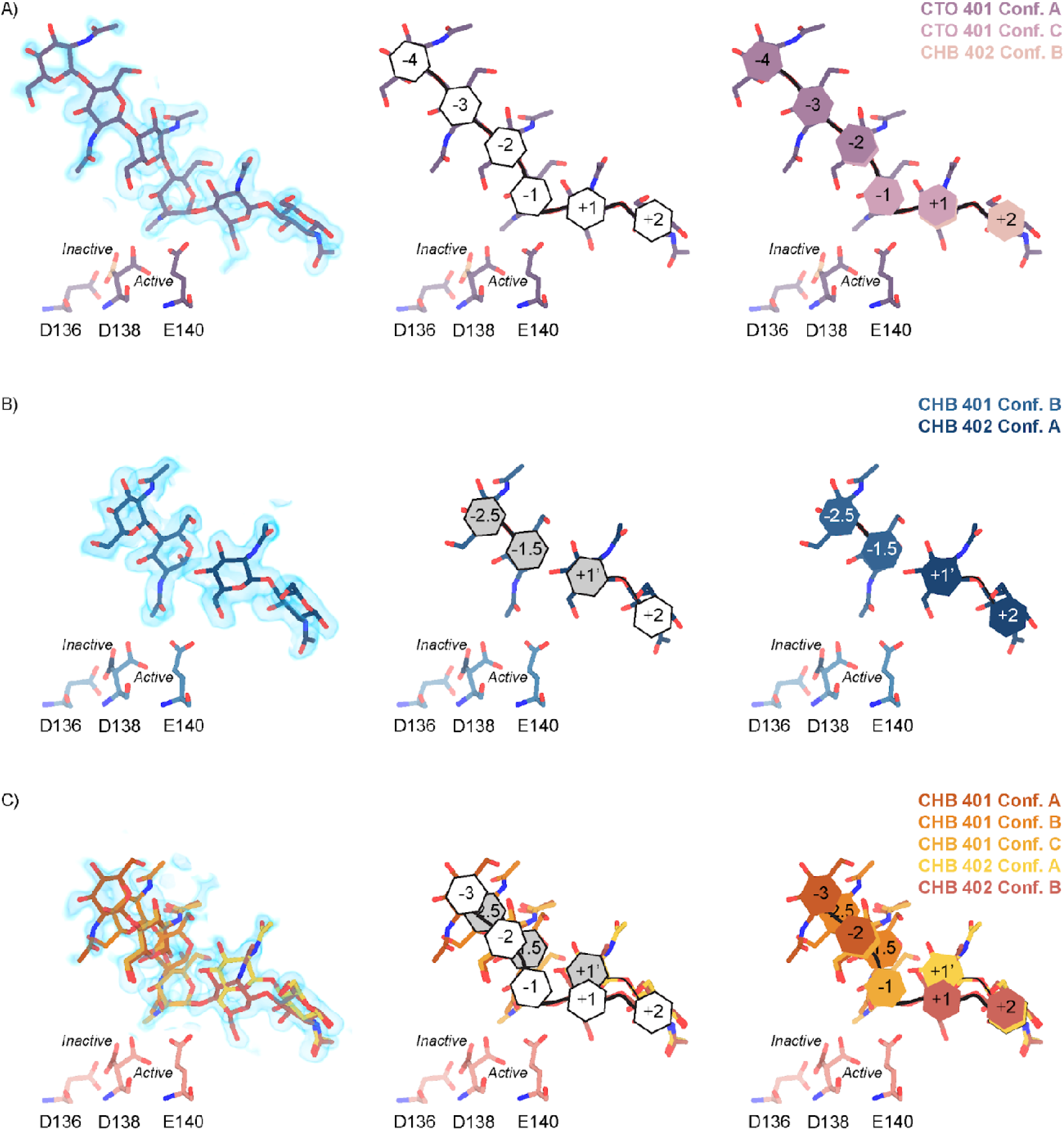
Schematic representation of sugar-binding subsites in mAMCase. **A)** PDB ID: 8GCA, chain A. Stick representation of all GlcNAc_2_ sugar-binding events observed in *n* sugar-binding subsites with 2mFo-DFc map shown as a 1.2 Å contour (blue), the subsite nomenclature, and a schematic of alternative conformation ligand modeling. **B)** PDB ID: 8FRA, chain D. Stick representation of all GlcNAc_n_ binding events observed in *n*+0.5 sugar-binding subsites with 2mFo-DFc map shown as a 1.2 Å contour (blue), the subsite nomenclature, and a schematic of alternative conformation ligand modeling. **C)** PDB ID: 8FR9, chain B. Stick representation of all GlcNAc_n_ binding events observed in *n* and *n*+0.5 sugar-binding subsites with 2mFo-DFc map shown as a 1.2 Å contour (blue), the subsite nomenclature, and a schematic of alternative conformation ligand modeling.

In addition to identifying novel *n*+0.5 sugar-binding subsites, we also observed strong positive difference density above the +1 subsite, which we label +1’. During ligand refinement, we observed density for both the α- and β-1,4-linked GlcNAc_2_ anomers in the active site. This unexpected configurational heterogeneity, which is observable because of the high resolution of our datasets (1.30 - 1.95 AU), likely formed as a result of equilibration between the two anomers through an oxocarbenium close-ion-pair intermediate. The ability for the active site to accommodate and form interactions with these ligands is important given its role in degrading crystalline chitin, a complex and often recalcitrant substrate that likely requires multiple binding events by AMCase before degradation can occur. We did not identify consistent trends between the contents of the crystallization drop (pH, substrate identity, and substrate concentration), the crystal properties (space group, unit cell dimensions, resolution), and the resulting density in the active site; however, as outlined below, the protein conformations and substrate states are highly correlated. Collectively, modeling a combination of ligand binding modes, linkages, and anomers allowed us to interpret the resulting coordinates in a more complete model of how mAMCase coordinates and stabilizes polymeric chitin for catalysis (**Figure 2**; **Supplemental Figure 5**D; **Supplemental Figure 6**A; **Table 2**).

**Table 2.** Occupancy of each ligand subsite and Asp138 in the active conformation (separate file).

### Structural characterization of mAMCase catalytic triad D_1_xD_2_xE

We interpreted the protein-ligand interactions along the canonical binding sites (**Supplemental Figure 5**C,D). As with other chitinases, we observe a network of tryptophans consisting of Trp31, Trp360, Trp99, and Trp218 stabilizing the positioning of the ligand into the binding site through a series of H-π interactions with the -3, -1, +1, and +2 sugars, respectively^25–27^. These interactions are primarily with the axial hydrogens of the respective sugars but also include the N-H of the -3 and +1 sugar and the 6’ O-H of the +2 sugar (**Supplemental Figure 6**B). Further, we observe Asp213 accepting a hydrogen bond with the 6’ OH of the -1 sugar and Tyr141 acting as a hydrogen bond donor to the 6’ OH of the +1 sugar. These two hydrogen bonds likely orient the ligand in the catalytically competent pose where the glycosidic oxygen bridging the -1 and +1 sugars is 2.8 AU away from the acidic Glu140 -OH (**Supplemental Figure 6**C). With this proximity, Glu140 can act as a hydrogen bond donor to the strained (122o bond angle) bridging oxygen forming a hydrogen bond to promote the formation of an oxazolinium intermediate and subsequent cleavage of the glycosidic bond. We observed two interactions with the sugar in the -4 position supporting the ligand orientation far from the enzymatic active site. Residues involved in ligand binding and catalysis adopt similar side chain conformations in the absence of ligand (PDB ID: 8FG5, 8FG7), suggesting that the active site is organized prior to ligand binding and not subject to ligand-stabilized conformational changes.

We hypothesize that the +1’ subsite is primarily occupied by the product GlcNAc_2_ prior to its displacement from the active site by subsequent sliding of polymeric chitin (**Figure 2**B)^28^. At this position, Trp99 and Trp218 engage in CH-π interactions with the +1 and +2 sugars, respectively while Asp213 forms a new H-bond with the carbonyl oxygen and Tyr141 retains an H-bond with the hydroxyl moiety on the +1 sugar. We are able to observe this post-catalysis binding mode due to the stabilizing interactions between GlcNAc_2_ and Asp213, Trp99, Trp218, and Tyr141 (**Supplemental Figure 6**C). Together, these observations highlight the dynamic chitin binding modes within the mAMCase active site. Collectively, the observed non-canonical binding modes of these sugars is consistent with previous observations that once bound to polymeric chitin, GH18 chitinases engage in chain sliding from the reducing end of the substrate following catalysis^29^.

In contrast to the largely static interactions outlined above, we observed conformational heterogeneity in the catalytically critical Asp138 residue, suggesting flipping between two equally stable states facing each of the other two residues in the catalytic triad (Asp136 or Glu140)^30^. Using Ringer, we confirmed that there are two Asp138 conformations and only a single conformation for Asp136 and Glu140 (**Supplemental Figure 7**; Data available at doi: 10.5281/zenodo.7758815)31. Across 20 chains from the datasets derived from different pH and co-crystallization conditions (**Table 1**), we quantified whether Asp138 is preferentially oriented towards Asp136 (*inactive* conformation) or preferentially oriented towards Glu140 (*active* conformation).

Prior work has suggested that Asp138 orients itself towards Glu140 to promote stabilization of the substrate’s twisted boat conformation in the -1 subsite. Therefore, we explored if Asp138 conformation is correlated with ligand pose^13,30,32,33^. As previously mentioned, we assign alternative conformation IDs to each ligand molecule based on its subsite positioning. We calculate subsite occupancy by taking the sum of all alternative ligand conformations at a given subsite, i.e. the occupancy of subsite -2 is equal to the occupancies of GlcNAc_2_ ResID 401 Conf. A and GlcNAc2 ResID 401 Conf. C (**Figure 3**A; see Methods for additional details; Data available at doi: 10.5281/zenodo.7905828). We observe a strong positive correlation between Asp138 conformation and ligand pose only in the -2 to +1 subsites (**Figure 3**B; **Table 2**). When the -1 subsite is at least 50% occupied, Asp138 prefers the *active* conformation (up towards Glu140). In this orientation, Asp138(HD2) forms a H-bond with Glu140(OE1) (2.6 AU) while Asp138(OD1) forms an H-bond with the amide nitrogen of GlcNAc in the -1 subsite (2.6 AU). Glu140(OE2) is 2.8 AU away from the glycosidic oxygen bridging the -1 and +1 sugars. We suspect that the inverse correlation between Asp138 *active* conformation and the -2.5 and -1.5 sugar-binding subsites represents ligand translocation towards the catalytic residues, prior to enzyme engagement with the ligand. When chitin occupies a canonical sugar-binding subsite, AMCase forms stabilizing H-bonds with the ligand prior to catalysis. These observations are consistent with the proposed catalytic mechanism where upon protonation, the equilibrium between Asp138 conformations shifts to favor the *active* conformation (towards Glu140) where Asp138 stabilizes Glu140 in proximity to the glycosidic oxygen prior to catalysis.

**Figure 3.**
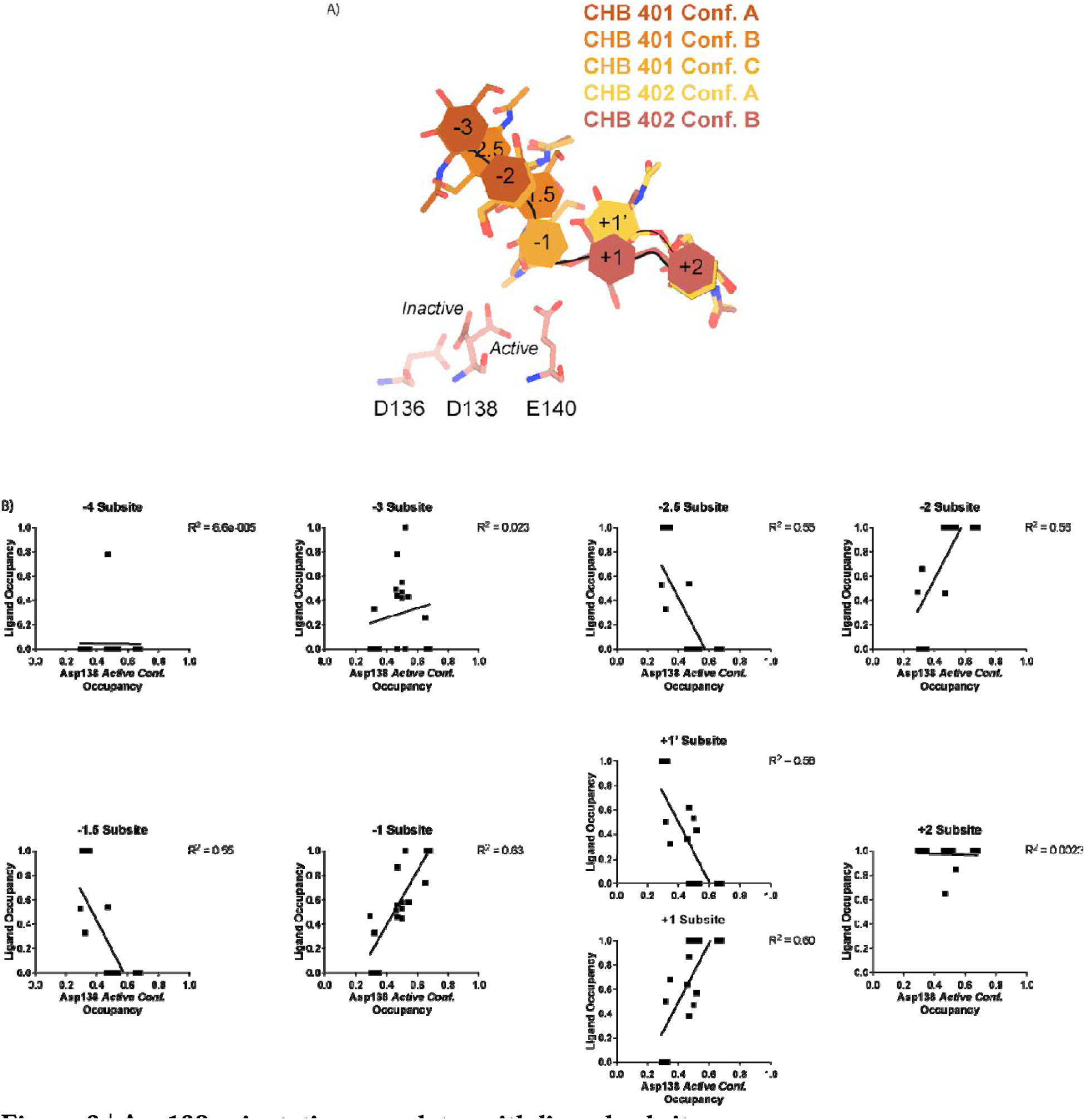
Asp138 orientation correlates with ligand subsite occupancy. **A)** PDB ID: 8FR9, chain B. Schematic of the alternative conformation ligand modeling. **B)** Linear correlation between sugar-binding subsite occupancy and Asp138 *active* conformation occupancy.

### Theoretical pKa calculations of mAMCase catalytic triad D_1_xD_2_xE

Based on the dual pH optimum observed in our kinetics assay and the conformational heterogeneity of Asp138, we calculated the theoretical pKa for catalytic D1xD2xE motif on mAMCase using PROPKA 3.0. PROPKA does not account for alternative conformations in its calculations, so we split our protein models to contain single conformations of the catalytic residues Asp136, Asp138, and Glu140. While PROPKA does account for ligands in its calculations, running the calculations with different alternative conformations of GlcNAc_2_ or GlcNAc_3_ had little effect on the calculated pKas for the active site residues (**Supplemental Figure 4**; Data available at doi: 10.5281/zenodo.7905863). Despite the observed ligand heterogeneity, we observe a relatively narrow range of pKa values for the catalytic triad. This suggests that the pKa of the catalytic residues is primarily influenced by the position of nearby residues and that the placement of solvent or ligand molecules has little effect. When Asp138 is oriented towards Asp136 (*the inactive* conformation), the pKa of the catalytic residues are 2.0, 13.0, 7.7 for Asp136, Asp138, and Glu140 respectively. Similarly, when Asp138 is oriented towards Glu140 (*the active* conformation), the pKa of the catalytic residues are 3.4, 12.4, 6.4 for Asp136, Asp138, and Glu140 respectively. Taking this information together, it is clear that the pKa of Asp136 and Glu140 are both affected by the orientation of Asp138 (**Figure 4**A; **Table 3**;

**Figure 4.**
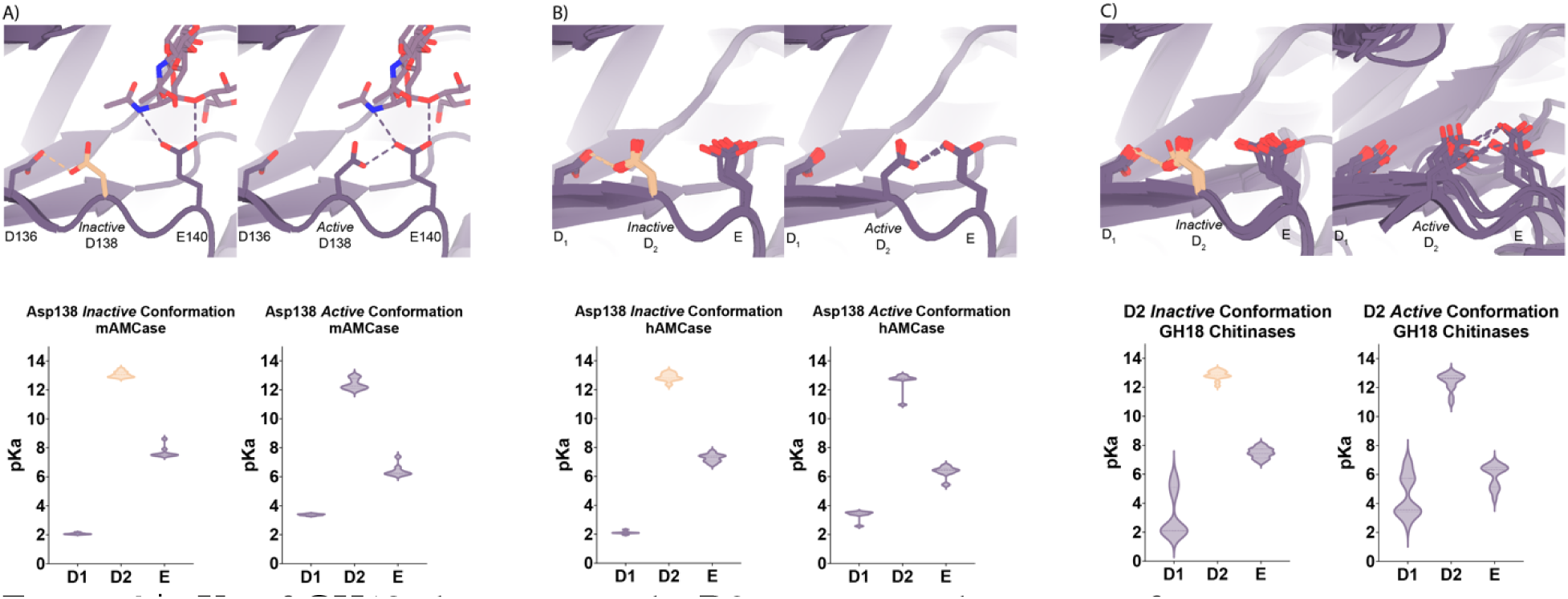
pKa of GH18 chitinases in the D2 inactive and active conformation. **A)** PDB ID: 8GCA, chain A. Distribution of pKa across Asp136, Asp138, Glu140 of mAMCase structures in either Asp138 *inactive* or Asp138 *active* conformation. **B)** PDB ID: 3FXY, 3RM4, 3RM8, 3RME (*inactive conformation)*; 2YBU, 3FY1 (*active* conformation). Distribution of pKa across Asp136, Asp138, Glu140 of hAMCase structures in either Asp138 *inactive* or Asp138 *active* conformation. **C)** PDB ID: 3ALF, 3AQU, 3FXY, 3RM4, 3RM8, 3RME (*inactive* conformation); 2UY2, 2UY3, 2YBU, 4HME, 4MNJ, 4R5E, 4TXE (*active* conformation). Distribution of pKa across the catalytic triad D_1_xD_2_xE of GH18 chitinases in either D_2_ *inactive* or *active* conformation.

**Table 3.** pKa across Asp136, Asp138, Glu140 of mAMCase structures in either Asp138 inactive or Asp138 active conformation (separate file).

Data available at doi: 10.5281/zenodo.7905863). The pKa of Asp136 suggests that at pH > 3.4, Asp136 is deprotonated, and its conjugate base is more stable. We observe a similar pKa distribution for the catalytic triad in human AMCase and other GH18 chitinases with publicly available structures and optimum pH activity profiles (**Figure 4**A-C). Given the pH range of our crystallization conditions, we expect that Asp136 is deprotonated while Asp138 and Glu140 are protonated. We hypothesize that this anionic aspartate is capable of forming a strong ionic hydrogen bond interaction with Asp138 orienting it in the *inactive* conformation. When Asp136 is protonated to its aspartic acid state at pH < 3.2, we expect that it is only capable of forming the relatively weaker neutral hydrogen bond with Asp138 lowering the favorability of the *inactive* conformation. Additionally, when interpreting the pKa of Glu140, we hypothesize that under acidic conditions (pH 2.0 - 6.5), Glu140 is capable of obtaining its catalytic proton from solution. The accessibility of Asp138’s proton to Glu140 progressively decreases as pH increases from pH 2.0 to 6.5. In contrast, under neutral and basic conditions (pH 6.0 - 7.4), Asp138 can shuttle a proton from Asp136 by rotating about its Cα-Cβ bond to supply Glu140 with the proton. Glu140 subsequently uses the proton that it obtained from Asp138 to protonate the glycosidic bond in chitin, promoting hydrolysis as previously described in several chitinases^30,34,35^. While this mechanism could explain how mAMCase has a local optimum at pH 2.0, it is insufficient to explain why we do not observe a similar optimum in hAMCase. The narrow range of pKa values across GH18 chitinases suggest that differences in optimal activity by pH may be influenced by other factors, such as protein stability, conformational dynamics, or coordination of distal GlcNAc residues by ionizable residues^36^.

### Molecular Dynamics

Based on our enzymology results suggesting the possibility of differential activity between acidic pH (pH 2.0) and near neutral pH (pH 6.5) and theoretical pKa calculations of the active site residues, we performed short atomistic molecular dynamics simulations to interrogate the movement of catalytic residues. While all the crystal structures we obtained were collected in a narrow acidic pH range between 4.74 - 5.60, we ran simulations at pH 2.0 and pH 6.5, ensuring that the protonation states of side chains populated by 3DProtonate were supported by our PROPKA calculations (Data available at doi: 10.5281/zenodo.7758821)37^,38^. These simulations allowed us to investigate our hypothesis that at neutral pH mAMCase enzymatic activity is dependent on the protonation state of Asp136. We performed simulations using protein models that contain Asp138 in either the *inactive* (down towards Asp136; “*inactive* simulation”) or *active* conformation (up towards Glu140; “*active* simulation”) to avoid bias from the starting conformation. In all our simulations, we observe that Glu140 orients its acidic proton towards the glycosidic bond between the -1 and +1 sugars. The distance between the acidic proton of Glu140 and the glycosidic oxygen fluctuates between 1.5 to 2.3 AU for the duration of the simulation, with a median distance of 1.8 AU. The positioning of this proton is necessary to allow for the oxocarbenium cleavage of the glycosidic bond and recapitulates the positioning of Glu140 in our experimental structures. In simulations initiated from the *inactive* conformation at pH 2.0, we observe that Asp 138 is readily able to rotate about its Cα-Cβ bond to adopt the *active* conformation forming the same hydrogen bond between Asp138 and Glu140. In contrast, from simulations at pH 6.5 started from the Asp138 *inactive* conformation, we observe that Asp138 remains hydrogen bonded to Asp136 throughout the duration of the simulation (*inactive* conformation; **Figure 5**A-C; Data available at doi: 10.5281/zenodo.7758821). This series of simulations allowed us to better visualize which catalytic side chains are dynamic and which catalytic side chains positioning are well maintained to help build our catalytic mechanism.

**Figure 5.**
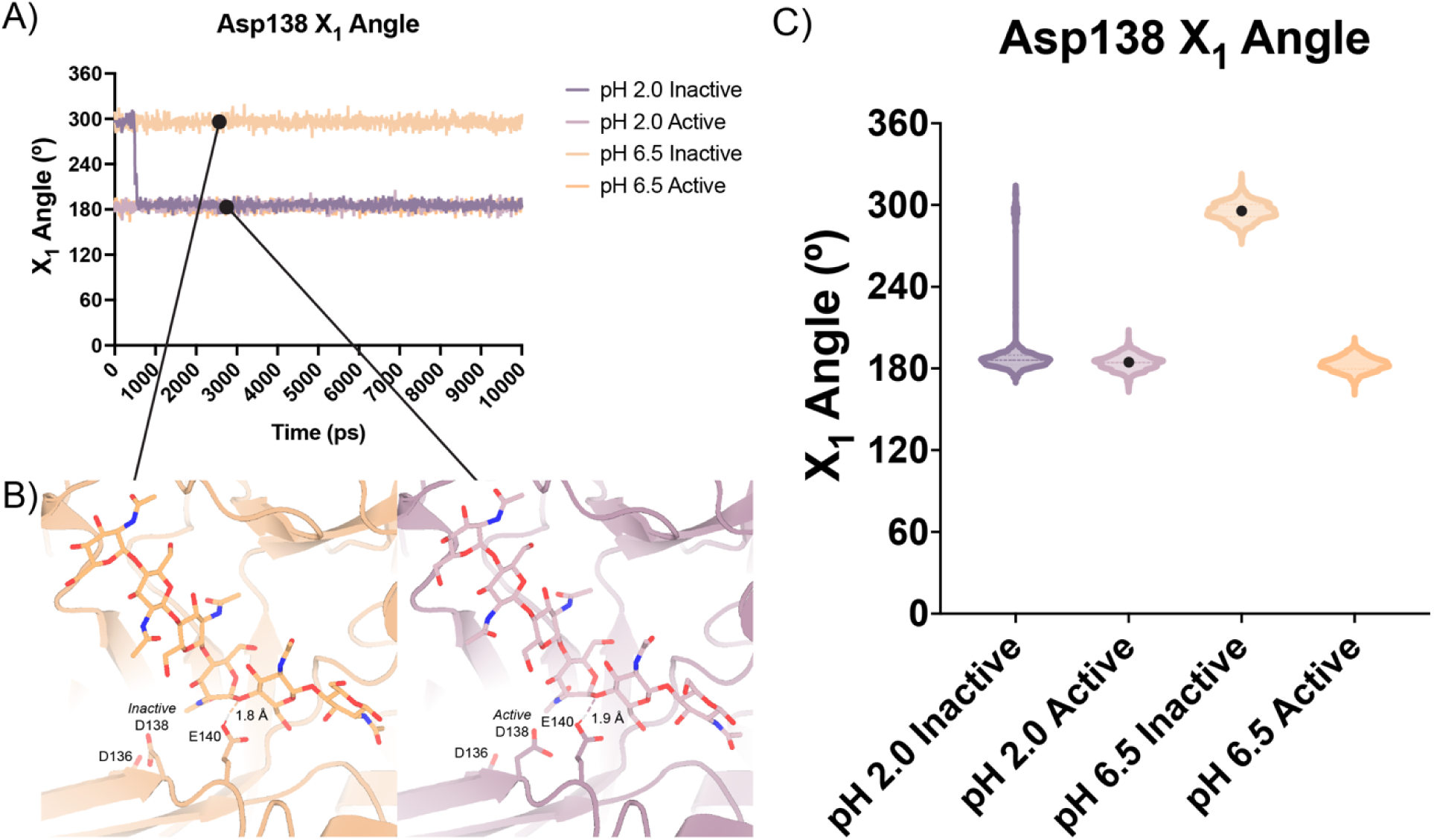
Distribution of distances observed every 10 ps of each simulation and their respective time courses. **A)** Asp138 χ_1_ angles over a 10 ns simulation. **B)** Representative minimum distance snapshots of structure during pH 6.5 *inactive* simulation (left), and pH 2.0 *active* simulation (right). **C)** Distribution of Asp138 χ_1_ angles over a 10 ns simulation.

## Discussion

mAMCase is an unusual enzyme that can bind and degrade polymeric chitin in very different pH environments. We hypothesized that mAMCase employs different mechanisms to protonate its catalytic glutamate under acidic and neutral pH. Through our analysis, we hypothesize that the observed ligand and catalytic residue densities and occupancies in our crystal structures are consistent with the previously proposed GH18 catalytic mechanism^39^. By modeling GlcNAc_2_ as sequentially overlapping ligands in alternative conformations (**Figure 2**), we are able to visualize each step in the proposed catalytic cycle of mAMCase (**Figure 6**). This mechanism, which has been observed in other glycoside hydrolases, occurs when the glycosidic oxygen is protonated by an acidic residue and a nucleophile adds into the anomeric carbon leading to elimination of the hydrolyzed product.

**Figure 6.**
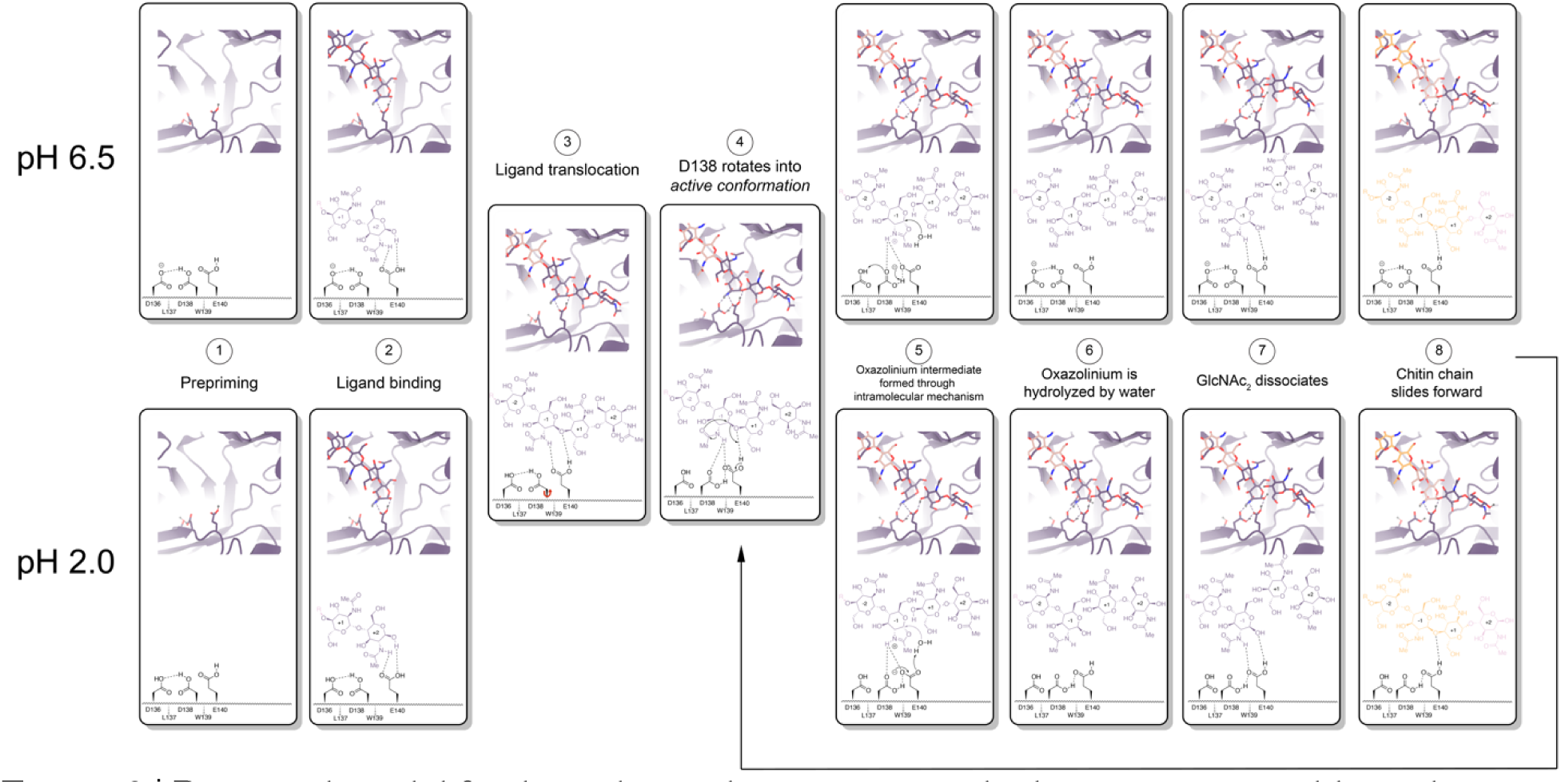
Proposed model for ligand translocation towards the active site and ligand release post-catalysis. **A)** PDB ID: 8GCA, chain A with no ligand (step 1); with GlcNAc_4_ generated by *phenix.elbow* using PubChem ID: 10985690 (step 2); with GlcNAc_6_ generated by *phenix.elbow* using PubChem ID: 6918014 (step 3-4, 8); with oxazolinium intermediate generated by *phenix.elbow* using PubChem ID: 25260046 (steps 5.1-5.2); with GlcNAc_2_ and GlcNAc_4_ generated by *phenix.elbow* using PubChem ID: 439544 and 10985690, respectively (steps 6-7). Chemical representation of GH18 catalytic cycle with corresponding molecular models of each step. Catalytic residues Asp136, Asp138, Glu140, and ligands are shown as sticks. Protons are shown as gray spheres. **B)** PDB ID: 8GCA, chain A. Animated movie of the mAMcase catalytic cycle at pH 2.0 (separate file) and **C)** at pH 6.5 (separate file). Catalytic residues Asp136, Asp138, Glu140, and ligands are shown as sticks. Protons are shown as gray spheres.

Based on our crystal data and simulations, we envision that at neutral pH, Asp136 is deprotonated (pKa = 2.1) forming an ionic hydrogen bond with Asp138 (pKa = 13.1). In contrast, at low pH Asp136 is protonated, yet continues to form a weaker hydrogen bond with Asp138 (**Figure 6** - Step 1). Glu140 (pKa = 7.7) is protonated across the enzyme’s active pH range. Upon ligand binding (**Figure 6** - Step 2), Glu140 stabilizes the sugar at the -1 subsite. The ligand then translocates forward by one GlcNAc_2_ to occupy the +1 and +2 subsites (**Figure 6** - Step 3). At neutral pH, Asp136 is predominantly deprotonated. When protonation of Asp136 occurs, this destabilizes the Asp136-Asp138 hydrogen bond and allows Asp138 to rotate about its Cα-Cβ bond into the *active* conformation (towards Glu140). However, since Asp136 is always protonated at low pH, the Asp136-Asp138 hydrogen bond is less energetically favorable, therefore Asp138 can adopt the *active* conformation more readily (**Figure 6**- Step 4).

Once Asp138 is in the *active* conformation, Asp138 and Glu140 form stabilizing interactions with the *N*-acetyl group of the ligand, priming it to become the nucleophile required for catalysis (**Figure 6** - Step 4). Glu140 provides its ionizable proton to the ligand’s glycosidic oxygen, increasing the electrophilicity of the anomeric carbon (**Figure 6** - Step 5)^40^. The carbonyl oxygen of the -1 sugar *N*-acetyl group then nucleophilically adds into the anomeric carbon from the β face to cleave the glycosidic bond, forming the oxazolinium intermediate. At neutral pH, the resultant deprotonated Glu140 is then re-protonated through proton shuttling in which Asp136 donates its proton to Asp138 and Asp138 donates its ionizable proton to Glu140. At acidic pH we propose that Glu140 can be directly re-protonated by a proton in solution (**Figure 6** - Step 5). At a neutral pH this leads to Asp138 returning to an *inactive* conformation. However, at low pH Asp136 and Glu140 are both protonated due to the high concentration of protons in solution, allowing Asp138 to remain in the *active* conformation and form stabilizing interactions with the *N*-acetyl group on the ligand. The oxazolinium intermediate is then hydrolyzed by a water molecule, generating a GlcNAc_2_ catalysis product in the +1 and +2 sugar subsites (**Figure 6** - Step 6). The GlcNAc_2_ product dissociates from the +1 to +2 sugar subsites, then the ligand undergoes “decrystallization” and “chain sliding” before re-entering the catalytic cycle, assuming AMCase is bound to a longer polymer such as its natural substrate^29^. At neutral pH this catalytic mechanism is reset with Asp138 in its *inactive* conformation, however at low pH the catalytic mechanism is reset with Asp138 already in the *active* conformation. This could lead to faster rates of catalysis at lower pH compared to the neutral pH mechanism, providing a possible explanation for the observed changes in rate at varying pH.

While our model helps us propose a plausible explanation of why mAMCase is highly active at pH 2, it does not explain why hAMCase has a single activity optimum around pH 5. Prior work by Kashimura et al. has demonstrated that *E. coli*-expressed mAMCase is remarkably stable across a broad pH range^41^. Similar experiments have not yet been performed on hAMCase. Olland et al. previously identified Arg145, His208, and His269 as important for pH specificity^13^. Seibold et al. argued that hAMCase isoforms containing asthma protective mutations N45D, D47N, and M61R, which are wildtype in mAMCase, may influence the pKa of Asp138-Glu140 by undergoing structural rearrangement^18^. Tabata et al. identified mutations across the course of evolution in Carnivora that were inactivating or structurally destabilizing (loss of S-S bonds)^42^. Okawa et al identified how primate AMCase lost activity by integration of specific, potentially pKa-shifting, mutations relative to the mouse counterpart^42b^.

To this end, we explored sequence differences between mouse and human AMCase homologs for insight into why mAMCase has such high enzymatic activity at pH 2.0 and 6.5 compared to hAMCase. We identified ionizable residues on mAMCase that likely contribute to its overall stability and are not present in hAMCase. Mutations Lys78Gln, Asp82Gly, and Lys160Gln result in the loss of surface-stabilizing salt bridges in hAMCase and may contribute to its reduced activity at more acidic pH. It is likely that the dual pH optima of mAMCase is intrinsic to the catalytic mechanism, where Glu140 can be protonated directly from solution (at low pH) or through proton shuttling across the catalytic triad (at neutral pH; **Figure 1**E). However, hAMCase is likely too destabilized at low pH to observe an increase in *k*_cat_. hAMCase may be under less pressure to maintain high activity at low pH due to humans’ noninsect-based diet, which contains less chitin compared to other mammals with primarily insect-based diets^42^. Together, these data demonstrate the importance of using structural and biochemical assays to develop our understanding of the catalytic mechanism governing mAMCase activity. Using biochemical and structural methods, we have developed a detailed model of how AMCase fulfills its role in chitin recognition and degradation. Small chitin oligomers are ideal for measuring the ability of AMCase to cleave β-1,4-glycosidic linkages between GlcNAc units, but these small oligomers do not represent the complex crystalline chitin encountered by AMCase in the lung. It is difficult to extrapolate the effects we observe using small chitin oligomers to binding (*k*on), processivity (*k*proc), catalysis (*k*cat), or product release (*k*off) on the native large and heterogeneous oligomeric substrates. In the future, we hope to be able to directly visualize the mAMCase-chitin interactions and characterize each step of the catalytic mechanism including decrystallization, degradation, product release, and chain sliding (also known as processivity).

To further understand the impact of pH on the structure of AMCase, it will be necessary to crystallize AMCase across a broader pH range that may expose conformational and structural changes that contribute to mAMCase’s unique pH activity profile. Our simulations have important limitations that could be overcome by quantum mechanical simulations that allow for changes in protonation state and improved consideration of polarizability. Further, neutron diffraction crystallography could provide novel critical insight into the placement of protons across the active site and help to develop a more complete model of mAMCase’s catalytic mechanism at different pH. Understanding the mechanistic basis behind an enzyme’s dual pH optima will enable us to engineer proteins with tunable pH optima to develop improved enzyme variants for therapeutic purposes for diseases, such as asthma and lung fibrosis.

## Supporting information

STable 3.2

STable 3.3

SFigure 3.6B

SFigure 3.6C

## Contributions

R.E. Díaz designed and cloned the construct that yielded the apo and holo mAMCase crystals; expressed, purified, and established crystallization conditions; collected X-ray diffraction data and processed data; modeled, refined, and analyzed structures; set up the 4MU-CB kinetic endpoint assay, collected, and analyzed data for the 4MU-CB assay; prepared the manuscript. A.K. Ecker designed the molecular dynamics simulation parameters, performed the simulations, and analyzed the data; developed the catalytic mechanism model for pH 2.0 and 6.5; prepared the manuscript. G.J. Correy vitrified crystals, collected X-ray diffraction data and processed data. P. Asthana collected X-ray diffraction data and processed data. I.D. Young processed the diffraction data. B. Faust assisted with mammalian cell culture for protein expression. I.B. Seiple provided access to the MOE software for MD simulations. S.J. Van Dyken guided the biochemistry work; prepared the manuscript. R. Locksley guided the biochemistry work; prepared the manuscript. M.C. Thompson guided the X-ray crystallography experiments, assisted with X-ray diffraction data processing and protein modeling. J.S. Fraser supervised work; prepared the manuscript; arranged funding.

## CRediT Author Statemente 1

Conceptualization, R.E.D., A.K.E., S.J.V.D., R.M.L., M.C.T., J.S.F. Methodology, R.E.D., A.K.E., B.F., M.C.T., J.S.F. Software, R.E.D. Investigation, R.E.D., A.K.E., G.J.C., P.A. Formal Analysis, R.E.D., A.K.E., G.J.C., I.D.Y., J.S.F. Visualization, R.E.D., A.K.E., J.S.F. Resources, B.F., I.B.S., J.S.F. Data Curation, R.E.D. Supervision, R.E.D., J.S.F. Project Administration, J.S.F. Writing – Original Draft, R.E.D., A.K.E., J.S.F. Writing – Review & Editing, R.E.D., A.K.E., S.J.V.D., R.M.L., J.S.F. Funding Acquisition, R.E.D., J.S.F.

## Data Availability

Raw experimental data, processing files, log files, GraphPad PRISM files, PyMOL scripts, and PyMOL sessions can be found on Zenodo or Protein Diffraction.

⌷Figure 1 | Kinetic properties of mAMCase catalytic domain at various pH.

◦doi: 10.5281/zenodo.8250616
⌷Figure 2 | Schematic representation of sugar-binding subsites in mAMCase.

◦doi: 10.5281/zenodo.7967930
⌷Figure 3 | Asp138 orientation correlates with ligand subsite occupancy

◦doi: 10.5281/zenodo.7905828
⌷Figure 4 | pKa of GH18 chitinases in the D2 inactive and active conformation.

◦doi: 10.5281/zenodo.7905863
⌷Figure 5 | Distribution of distances observed every 10 ps of each simulation and their respective time courses.

◦doi: 10.5281/zenodo.7758821
⌷Figure 6 | Proposed model for ligand translocation towards the active site and ligand release post-catalysis.

◦doi: 10.5281/zenodo.7967958
⌷Supplemental Figure 1 | pH of reaction solution before and after quenching with 0.1 M Gly-NaOH pH 10.7

◦doi: 10.5281/zenodo.8250616
⌷Supplemental Figure 2 | Kinetics of 4MU-chitobioside catalysis by mAMCase catalytic domain at various pH.

◦doi: 10.5281/zenodo.8250616
⌷Supplemental Figure 3 | 96-well plate layout of crystallization conditions.

◦doi: 10.5281/zenodo.7905944
◦doi: 10.18430/M38FG5
◦doi: 10.18430/M38FG7
◦doi: 10.18430/M38GCA
◦doi: 10.18430/M38FRC
◦doi: 10.18430/M38FR9
◦doi: 10.18430/M38FRB
◦doi: 10.18430/M38FRD
◦doi: 10.18430/M38FRG
◦doi: 10.18430/M38FRA
⌷Supplemental Figure 4 | pKa of apo and holo mAMCase in the D2 *inactive* and *active* conformation.

◦doi: 10.5281/zenodo.7905863
⌷Supplemental Figure 5 | Overview of key residues for mAMCase activity.

◦doi: 10.5281/zenodo.7967978
⌷Supplemental Figure 6 | Protein-ligand interactions between mAMCase and chitin.

◦doi: 10.5281/zenodo.7967954
⌷Supplemental Figure 7 | Ringer analysis of catalytic triad confirms alternative Asp138 conformations.

◦doi: 10.5281/zenodo.7758815

## Methods

### Protein expression and purification

Protein expression and purification mAMCase catalytic domain (UniProt: Q91XA9; residues 22 to 391) was cloned into a pTwist CMV [pmRED006; Twist Biosciences; Addgene ID: 200228] or pcDNA3.1(+) [pmRED013; Genscript; Addgene ID: 200229] expression vector with a C-terminal 6xHis tag. To express mAMCase catalytic domain, 0.8-1 µg/mL plasmid DNA was transfected into ExpiCHO-S cells (ThermoFisher Scientific #A29127) using the Max Titer protocol (ThermoFisher Scientific MAN0014337). After cells were grown shaking at 37°C with 8% CO2 for 18-22 hours, ExpiFectamine CHO Enhancer (ThermoFisher Scientific #A29129) and ExpiCHO feed (ThermoFisher Scientific #A29129) was added to the flask. Cells were then transferred to 32 °C with 5% CO2 for an additional 9-13 days of growth, with a second volume of ExpiCHO feed added to the flask on day 5 post-transfection. Cells were removed by centrifugation at 4,000 RCF for 15 minutes at 4 °C, and the remaining supernatant was filtered using a 0.22 µm filter at 4 °C. Filtered supernatant was either dialyzed into Ni–nitrilotriacetic acid (NTA) loading buffer [100 mM Tris-HCl (pH 8.0), 150 mM NaCl] at 4 °C in a 10-kDa molecular weight cutoff (MWCO) Slide-A-Lyzer Dialysis Cassette, (ThermoFisher Scientific #66810) for 18-24 hours or concentrated in a 10-kDa MWCO centrifugal concentrator (Amicon #UFC901008) at 4,000 RCF in 5 min intervals until the final volume was equal to 10 mL, which was then diluted 1:10 with loading buffer for a total volume of 100 mL. The dialyzed supernatant volume was filtered using a 0.22 µm filter at 4 °C. All purification steps were performed at 4°C using an ÄKTA fast protein liquid chromatography system (Cytiva). The dialyzed supernatant was applied to a 5-ml HisTrap FF column (Cytiva, 17525501). The column was washed with 40 mL of loading buffer followed by 25 mL of 10% Ni-NTA elution buffer [100 mM Tris-HCl (pH 8.0), 150 mM NaCl, 500 mM imidazole] and then eluted over a 50 mL gradient from 10% to 100% elution buffer. Eluted protein was concentrated to 2.5 mL using a 10-kDa MWCO centrifugal concentrator (Amicon, UFC901024). The sample was further purified by size exclusion chromatography (SEC) using a HiLoad 16/600 Superdex 75 pg column (Cytiva, 28989333) equilibrated with SEC buffer [25 mM Tris-HCl (pH 8.0), 50 mM NaCl]. Eluted fractions were collected and stored at 4 °C for further use.

### 4MU-chitobioside Endpoint Assay

Chitinase catalytic activity has previously been assayed using 4-methylumbelliferyl chitobioside (4MU-CB; Sigma-Aldrich M9763) ^43,44^. 100 nM chitinase enzyme was incubated with varying concentrations of 4MU-chitobioside up to 117 μM in McIlvaine Buffer at 37 °C ^21^. The 4-methylumbelliferone (4MU) fluorophore is quenched by a ß-glycosidic linkage to a short chitin oligomer, which is cleaved by a chitinase enzyme, which generates fluorescence with peak excitation at 360 nm and emission at 450 nm. 4MU fluorescence is pH-dependent with peak excitation at 360 nm and emission at 450 nm at pH 7.0. It has been previously reported that 4MU peak excitation/emission increases and fluorescence intensity decreases as pH becomes more acidic ^45^. Given the pH-dependent fluorescence properties of the 4MU fluorophore, we incubate the reaction at different pH, then quench with 0.1 M Gly-NaOH pH 10.7. Quenching the reaction with 0.1 M Gly-NaOH pH 10.7 stops the enzyme reaction and shifts the pH to maximize the quantum yield of the 4MU substrate.

A Tecan Spark multimode microplate reader is pre-heated to 37 °C. 4MU-chitobioside (Sigma-Aldrich M9763) and AMCase are separately pre-incubated at 37 °C for 15 minutes. 25 µL of 4MU-chitobioside or McIlvaine Buffer (Boston Bioproducts) is transferred into each well in a Multiplate 96-Well PCR Plate, high profile, unskirted, clear (Bio-Rad MLP9601). Using a Multidrop Combi Reagent Dispenser (Thermo Scientific #5840300), 25 µL of either 100 nM AMCase or McIlvaine Buffer (Boston Bioproducts) is dispensed into each well in the Multiplate 96-Well PCR Plate (Corning #3993). The Multiplate 96-Well PCR Plate is then incubated at 37 °C in a 96-well Non-Skirted PCR Plate Block (Thermo Scientific #88870120) in a digital dry bath (Thermo Scientific #88870006).

The reaction is quenched with 50 µL 0.1 M Gly-NaOH pH 10.7 at timepoints 0”, 15”, 30”, 45”, 60”, 90”. 40 µL of the quenched reaction is transferred to a 384-well Low Volume Black Flat Bottom Polystyrene NBS Microplate (Corning #3820), then immediately read using the following parameters:

- Excitation - 360 nm, 20 nm bandwidth
- Emission - 450 nm, 20 nm bandwidth
- Gain - 50
- Flashes - 20

This assay was performed in quadruplicate for each pH unit reported. This allowed us to reliably measure initial rates of catalysis across a large range of pH conditions. The workflow for this assay is illustrated in **Figure 1**. A detailed protocol for this assay can be found on <u>protocols.io</u>.

### Analysis of kinetic data

25 µL of 200 µM 4MU fluorophore (Sigma-Aldrich M1381) was serially diluted into 25 µL McIlvaine Buffer (Boston Bioproducts) across the range of pHs to obtain 5 diluted ligand concentrations ranging from 100 µM to 6.25 µM as well as ligand free. This dilution series was performed in duplicate per 96-Well PCR plate for a total of 8 replicates per ligand concentration at each given pH value. At the end of the experiment, the 4MU dilution series is quenched with 50 µL 0.1 M Gly-NaOH pH 10.7 for a final dilution series ranging from 50 µM to 3.125 µM.

Relative fluorescence (RFU) was plotted against 4MU concentration, then a simple linear regression with the constraint Y = 0 when X = 0 was performed to obtain a standard curve. We then used the equation Y = mX + b, where m is the slope from the standard curve and Y is the RFU from a given experimental data point, to determine the concentration of 4MU [µM] generated by AMCase at a given time point.

Average 4MU concentration [µM] (n = 4) was plotted as a function of time with error bars representing the standard deviation. We then fit a simple linear regression with the constraint Y = 0 when X = 0 to obtain the initial rate of enzyme activity (4MU [µM]/sec) at each concentration of 4MU-chitobioside [µM]. Average initial rate (n = 4) was plotted as a function of 4MU-chitobioside concentration [µM] with error bars representing the standard deviation. We fit our data to a Michaelis-Menten function without substrate inhibition to obtain V_max_ and *K*_M_ parameters. We used the equation *k*_cat_ = V_max_/[Enzyme] where [Enzyme] = 0.1 µM to calculate *k*_cat_. We calculated catalytic efficiency (CE) using the equation CE = *K*_M_/*k*_cat_. Kinetic parameters Vmax, *K*_M_, *k*_cat_, and catalytic efficiency were plotted as a function of pH.

### Apo crystallization

Using hanging-drop vapor diffusion, crystallization screens were performed using a 96-well Clear Flat Bottom Polystyrene High Binding microplate (Corning CLS9018BC) with 0.5 mL of reservoir solution in each well. Crystallization drops were set up on 96-well plate seals (SPT Labtech 4150-05100) with 0.2 µl of AMCase at 11 mg/ml and 0.2 µl of reservoir using an SPT Labtech mosquito crystal. After 21 days at 20 °C, we observed crystals in a reservoir solution containing 20% PEG-6000 , 0.1 M Sodium Acetate pH 5.0, and 0.2 M Magnesium Chloride (II) (MgCl2) (NeXtal PACT Suite Well A10; #130718).

### Apo data collection, processing, and refinement at cryogenic temperature

Diffraction data were collected at the beamline ALS 8.3.1 at 100 K. Diffraction data from multiple crystals were merged using xia2^46^, implementing DIALS^47^ for indexing and integration, and Aimless^48^ for scaling and merging. We confirmed the space group assignment using DIMPLE ^49^. We calculated phases by the method of molecular replacement, using the program Phaser^50^ and a previous structure of hAMCase (PDB: 3FXY) as the search model. The model was manually adjusted in Coot to fit the electron density map calculated from molecular replacement, followed by automated refinement of coordinates, atomic displacement parameters, and occupancies using phenix.refine^51^ with optimization of restraint weights. Default refinement parameters were used, except the fact that five refinement macrocycles were carried out per iteration and water molecules were automatically added to peaks in the 2mFo-DFc electron density map higher than 3.5 Å. The minimum model-water distance was set to 1.8 Å, and a maximum model-water distance was set to 6 Å. For later rounds of refinement, hydrogens were added to riding positions using *phenix.ready_set*, and B-factors were refined anisotropically for non-hydrogen and non-water atoms. Following two initial rounds of iterative model building and refinement using the aforementioned strategy, we began introducing additional parameters into the model, enabled by the extraordinarily high resolution of our diffraction data. First, we implemented anisotropic atomic displacement parameters for heavy atoms (C, N, O, and S), followed by refinement of explicit hydrogen atom positions. A final round of refinement was performed without updating water molecules.

### Apo data collection, processing, and refinement at room temperature

Diffraction data were collected at the beamline ALS 8.3.1 at 277 K. Data collection, processing, refinement, and model building were performed as described previously for the apo crystals at cryogenic temperature.

### Holo crystallization

Initially, crystals were grown by hanging-drop vapor diffusion with a reservoir solution containing 20% PEG-6000 (Hampton Research HR2533), 0.1 M Sodium Acetate (pH 3.6, Hampton Research HR293301; pH 4.1, Hampton Research HR293306; pH 5.0, Hampton Research HR293315; pH 5.6, Hampton Research HR293321), and 0.2 M Magnesium Chloride (II) (MgCl2) (Hampton Research HR2559). Screens were performed using a 96-well Clear Flat Bottom Polystyrene High Binding microplate (Corning CLS9018BC) with 0.5 mL of reservoir solution in each well. Crystallization drops were set up on 96-well plate seals (SPT Labtech 4150-05100) with 0.2 µl of AMCase at 11 mg/ml and 0.2 µl of reservoir using an SPT Labtech mosquito crystal. Crystals grew after 1-2 days at 20 °C.

Using hanging drop diffusion vapor, holo crystals grew after 12 hours at 20 °C. For the holo form with GlcNAc_2_ (Megazyme O-CHI2), this construct crystallized in either P2_1_2_1_2 or P2_1_2_1_2_1_ with either 2 or 4 molecules in the ASU and diffracted to a maximum resolution between 1.50 to 1.95 Å. For the holo form with GlcNAc_3_ (Megazyme O-CHI3), this construct crystallized in P2_1_2_1_2 with 2 molecules in the ASU and diffracted to a maximum resolution of 1.70 Å.

### Holo data collection, processing, and refinement at cryogenic temperature

Diffraction data were collected at the beamline ALS 8.3.1 and SSRL beamline 12-1 at 100 K. Data collection, processing, refinement, and model building were performed as described previously for the apo crystals.

Ligands were modeled into 2mFo-DFc maps with Coot, using restraints generated by *phenix.elbow* from an isomeric SMILES (simplified molecular input line-entry system) string ^52^ using AM1 geometry optimization. Default refinement parameters were used, except the fact that five refinement macrocycles were carried out per iteration and water molecules were automatically added to peaks in the 2mFo-DFc electron density map higher than 3.5 Å. The minimum model-water distance was set to 1.8 Å, and a maximum model-water distance was set to 6 Å. Changes in protein conformation and solvation were also modeled. Hydrogens were added with *phenix.ready_set*, and waters were updated automatically. A final round of refinement was performed without updating water molecules^53^.

### Ligand modeling

For consistency, ligands were assigned an alternative conformation ID based on the sugar-binding subsites it occupied:

GlcNAc_2_ ResID 401 Conf. A, -3 to -2
GlcNAc_2_ ResID 401 Conf. B, -2.5 to -1.5
GlcNAc_2_ ResID 401 Conf. C, -2 to -1
GlcNAc_2_ ResID 402 Conf. D, -1 to +1
GlcNAc_2_ ResID 402 Conf. B, +1 to +2
GlcNAc_2_ ResID 402 Conf. A, +1’ to +2
GlcNAc_3_ ResID 401 Conf. A, -4 to -2
GlcNAc_3_ ResID 401 Conf. B, -3 to -1
GlcNAc_3_ ResID 401 Conf. C, -2 to +1
GlcNAc_3_ ResID 402 Conf. B, -1 to +2

Ligand occupancies and B-factors using *phenix.refine*. Ligands with occupancies <= 0.10 were removed from the model.

### Ringer analysis

Individual residues in each of the mAMCase structures were run through Ringer using mmtbx.ringer. Outputs from the csv file were then plotted using Matplotlib.

### pKa Analysis

We used the APBS-PDB2PQR software suite (https://server.poissonboltzmann.org/pdb2pqr)^54^. Each PDB model was separated into two separate models containing a single Asp138 conformation in either the *inactive* (down towards Asp136) or *active conformation* (up towards Glu140). Solvent and ligand molecules were not modified. The pH of the crystallization condition was provided for PROPKA to assign protonation states. The default forcefield PARSE was used. The following additional options were selected: Ensure that new atoms are not rebuilt too close to existing atoms; Optimize the hydrogen bonding network.

### Molecular Dynamics

Simulations were performed using hexaacetyl-chitohexaose (PubChem Compound ID: 6918014) modeled into 8GCA with Asp138 in either the *inactive* (down towards Asp136) or *active conformation* (up towards Glu140). The model PDB file was opened in MOE and solvated in a sphere of water 10 Å away from the protein. This system then underwent structural preparation for simulations using the standard parameters with the AMBER14 forcefield. The system then was protonated to set pH {2.0, 6.5} based on sidechain pKa predictions using the 3DProtonate menu followed by confirmation of appropriate protonation by PROPKA calculations. Protonated models underwent energy minimization by steepest descent before simulations were set up. Equilibration was performed for 10 ps followed by 100 ps of thermal gradient equilibration from 0K to 300K. A thermal bath equilibration was run for 100 ps before the production runs were started. Productions were run for 10 ns with a time step of 0.5 fs to not overshoot bond vibrations. The simulation was sampled every 10 ps for subsequent data analysis which was performed using the MOE database viewer and replotted using GraphPad Prism.

## Supplemental Figures

**Supplemental Figure 1.**
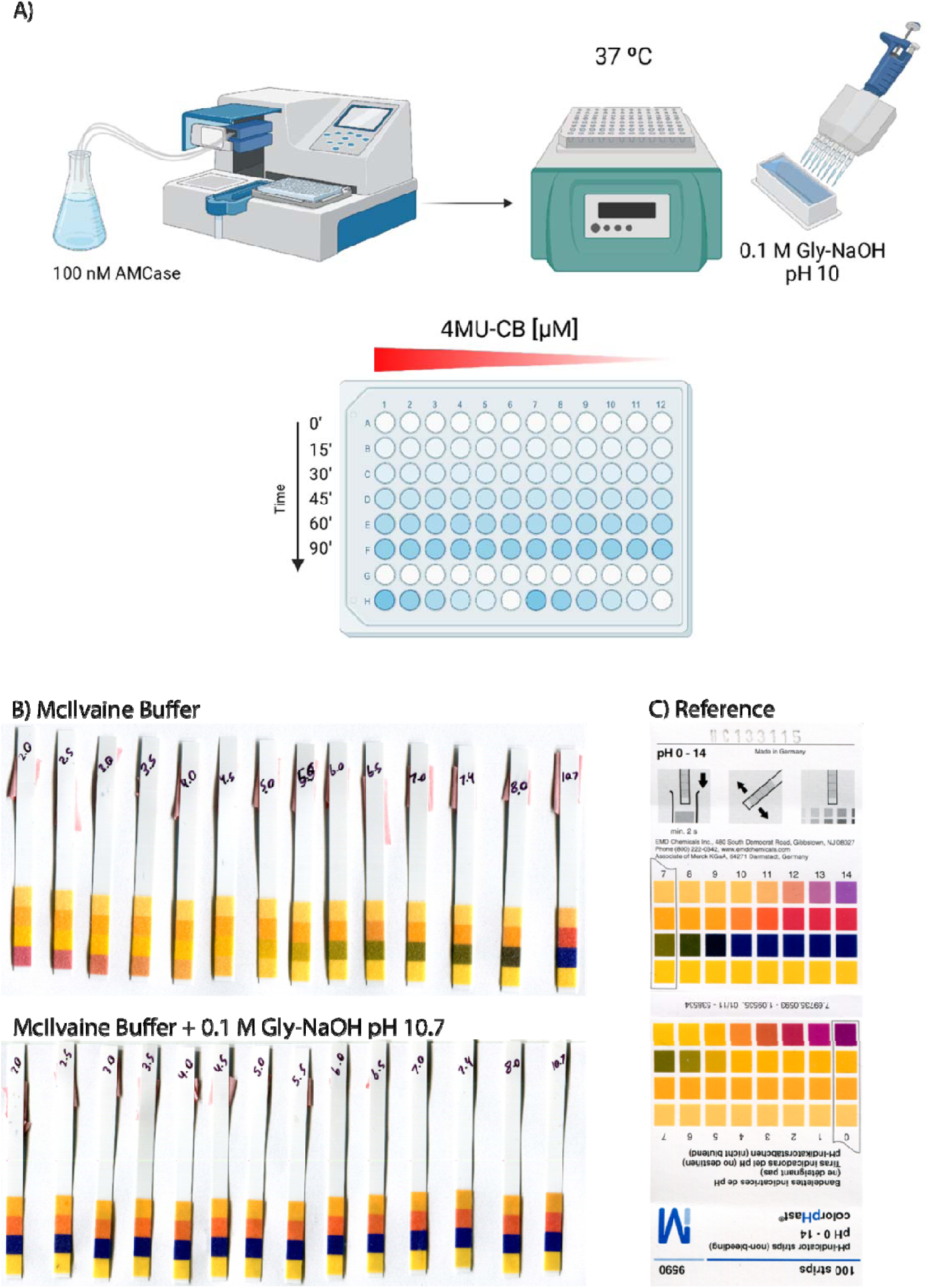
pH of reaction solution before and after quenching with 0.1 M Gly-NaOH pH 10.7. **A)** Schematic of modified endpoint 4MU-chitobioside assay. **B)** Reaction pH before and after quenching with 0.1 M Gly-NaOH pH 10.7, and **C)** a pH strip reference sheet.

**Supplemental Figure 2.**
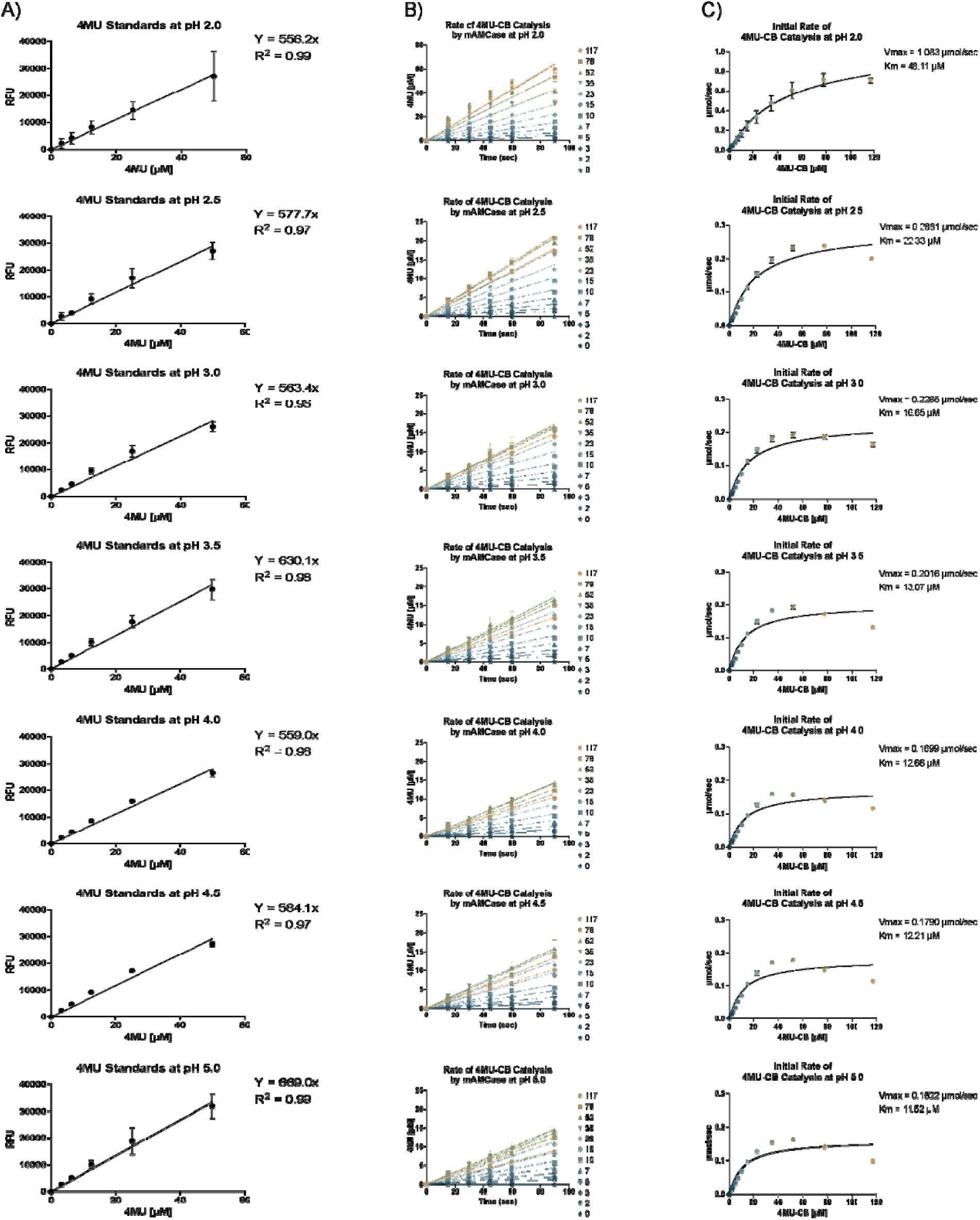
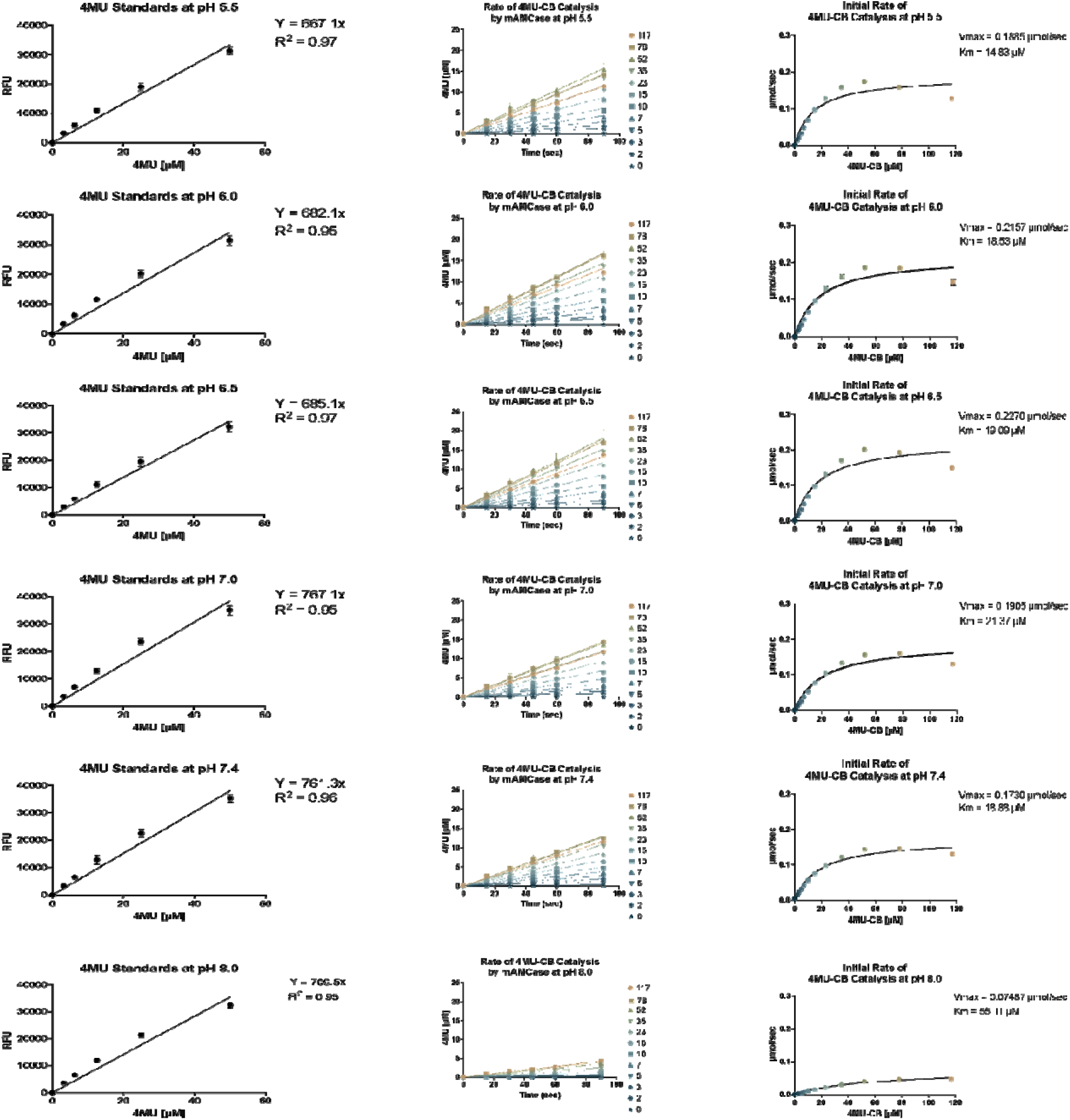
Kinetics of 4MU-chitobioside catalysis by mAMCase catalytic domain at various pH. **A)** A linear fit forced through Y = 0 is used to generate the standard curve for converting RFU to 4MU [µM]. Each data point represents n = 8 with error bars representing the standard deviation. **B)** 4MU fluorescence (RFU) is plotted as a function of time (sec). Each data point represents n = 4 with error bars representing the standard deviation. A linear fit is applied to each concentration of 4MU-chitobioside to calculate an initial rate. RFU is converted to µM using a 4MU standard curve. **C)** The rate of 4MU-chitobioside catalysis (1/sec) by mAMCase catalytic domain is plotted as a function of 4MU-chitobioside concentration (µM). Each data point represents n = 4 with error bars representing the standard deviation. Michaelis-Menten equation without substrate inhibition was used to estimate the *k*_cat_ and *K*_M_ from the initial rate of reaction at various substrate concentrations.

**Supplemental Figure 3.**
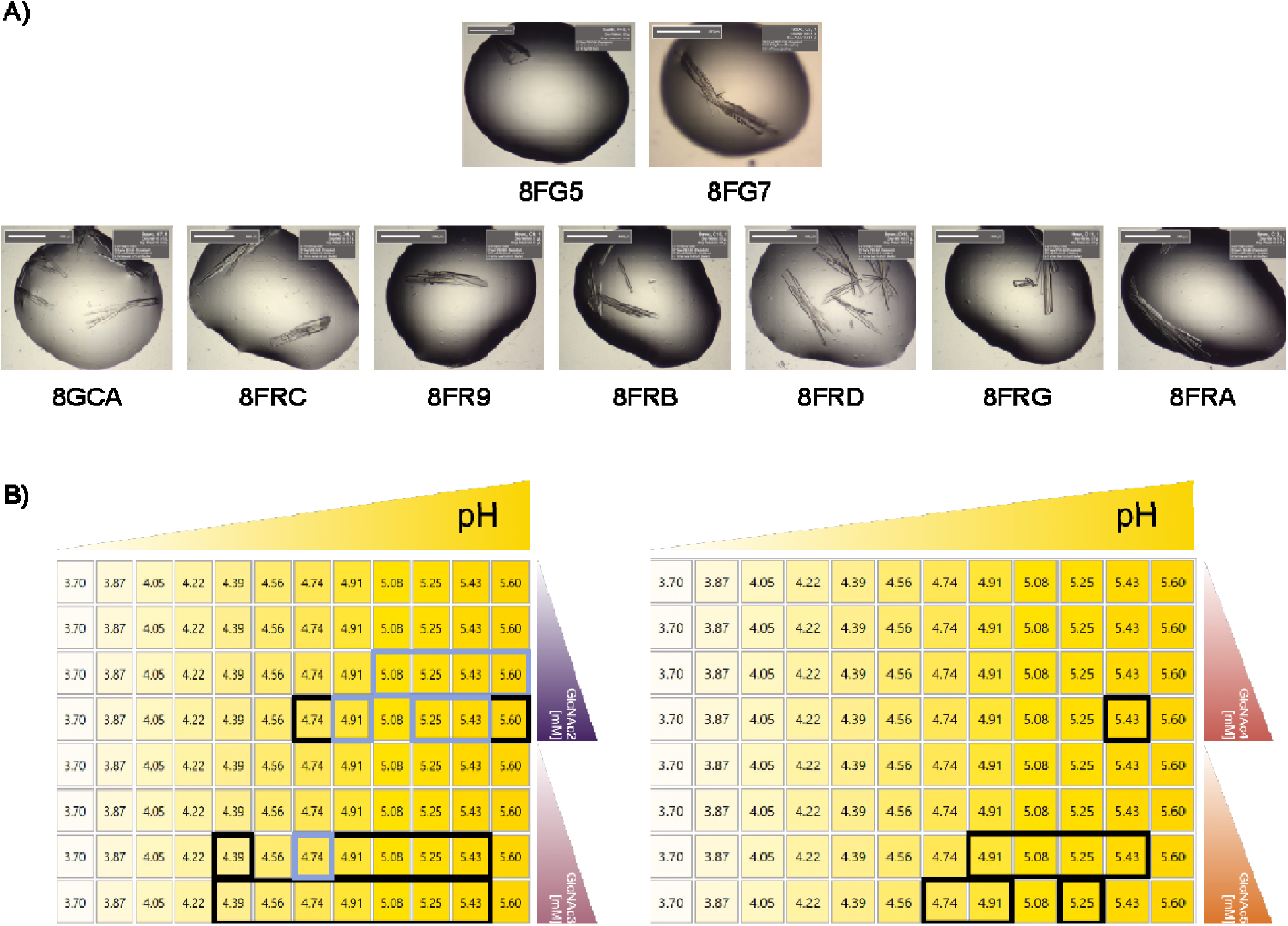
96-well plate layout of crystallization conditions. **A)** Brightfield view of crystals used to determine the structures reported in this paper. **B)** Hanging drop crystallization trays were set up as a 2-condition gradient to identify optimal crystallization conditions for AMCase + GlcNAc_n_. pH increased along the X-axis from pH 3.70 to 5.60. Ligand concentration increased along the Y-axis from 0 mM to 29 mM [GlcNAc_2_], 19 mM [GlcNAc_3_], 10 mM [GlcNAc_4_], or 8 mM [GlcNAc_5_]. Black boxes indicate conditions where crystals grew. Lilac boxes indicate conditions for structures reported in this paper.

**Supplemental Figure 4.**
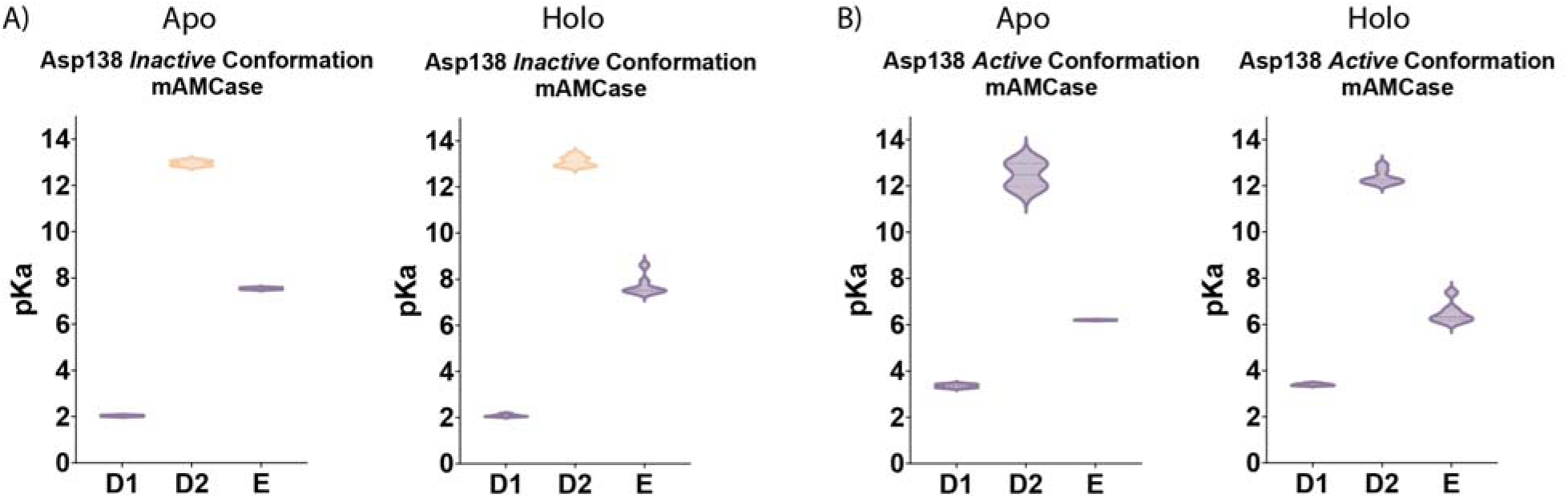
pKa of apo and holo mAMCase in the D2 *inactive* and *active* conformation. PDB ID: 8FG5, 8FG7 (*apo*); 8GCA, 8FRC, 8FR9, 8FRB, 8FRD, 8FRG, 8FRA (*holo*). Violin plots showing the distribution of pKa across Asp136, Asp138, Glu140 between **A)** apo and **B)** holo mAMCase structures in the *inactive* or *active* conformation.

**Supplemental Figure 5.**
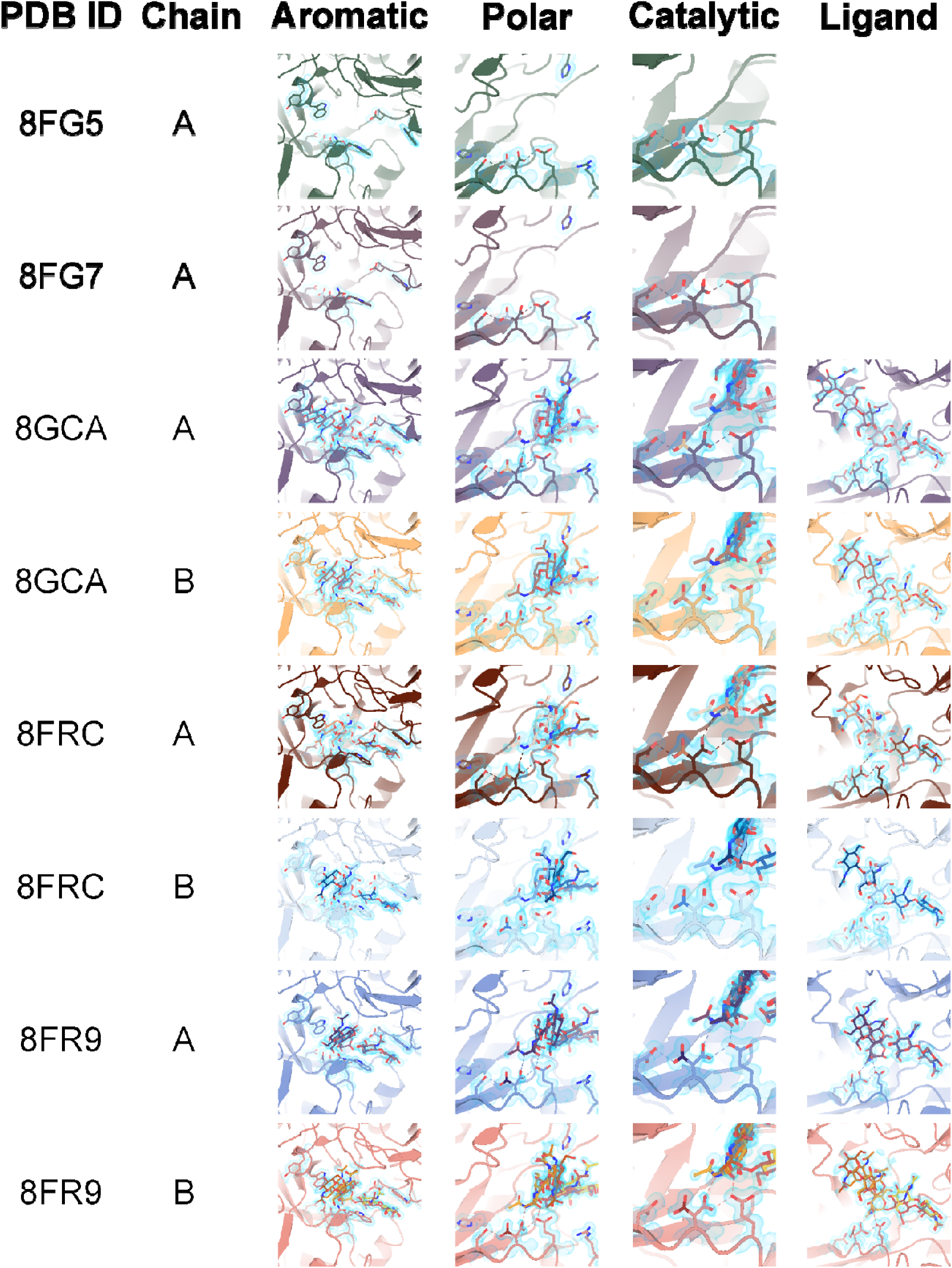
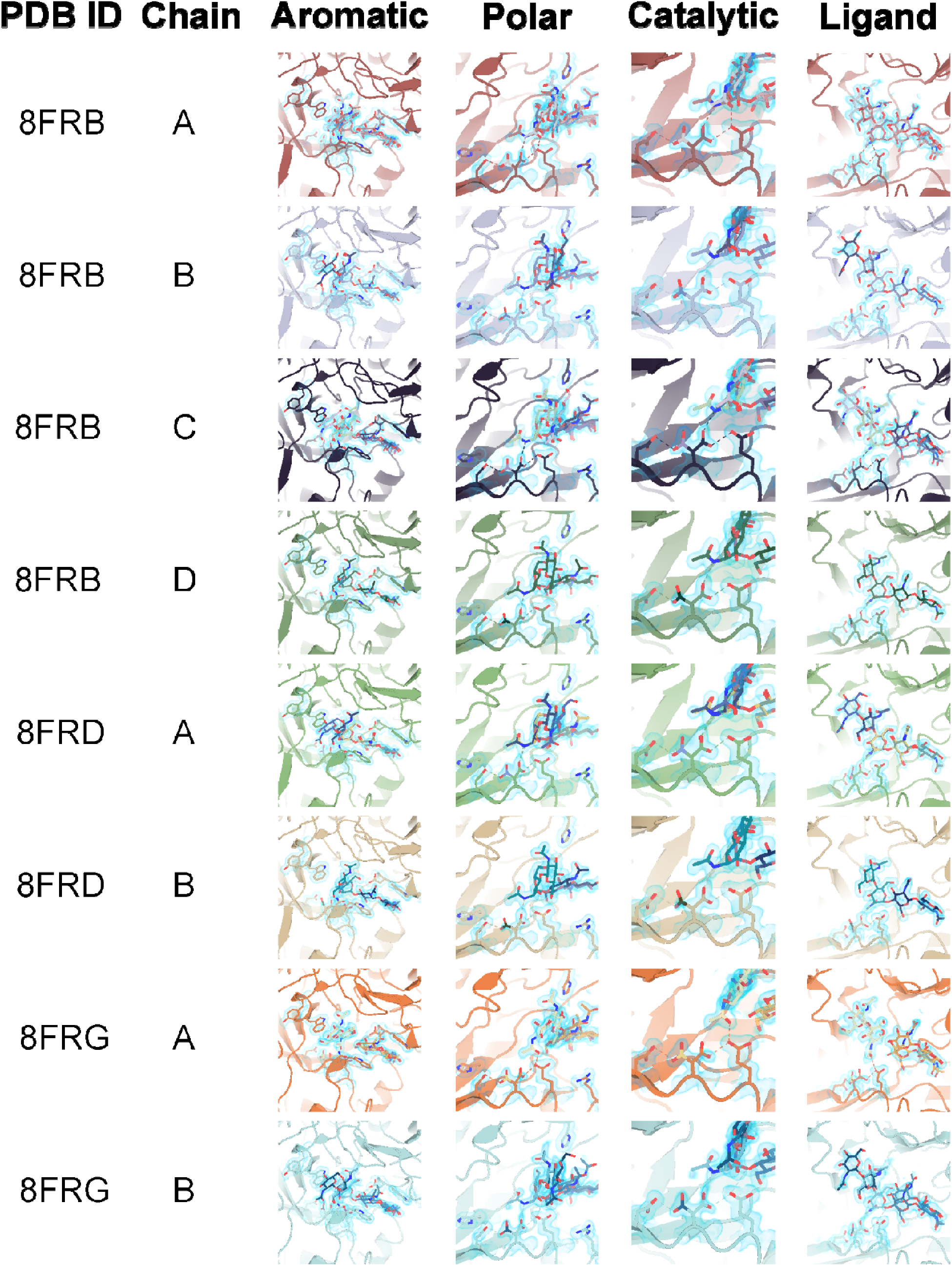
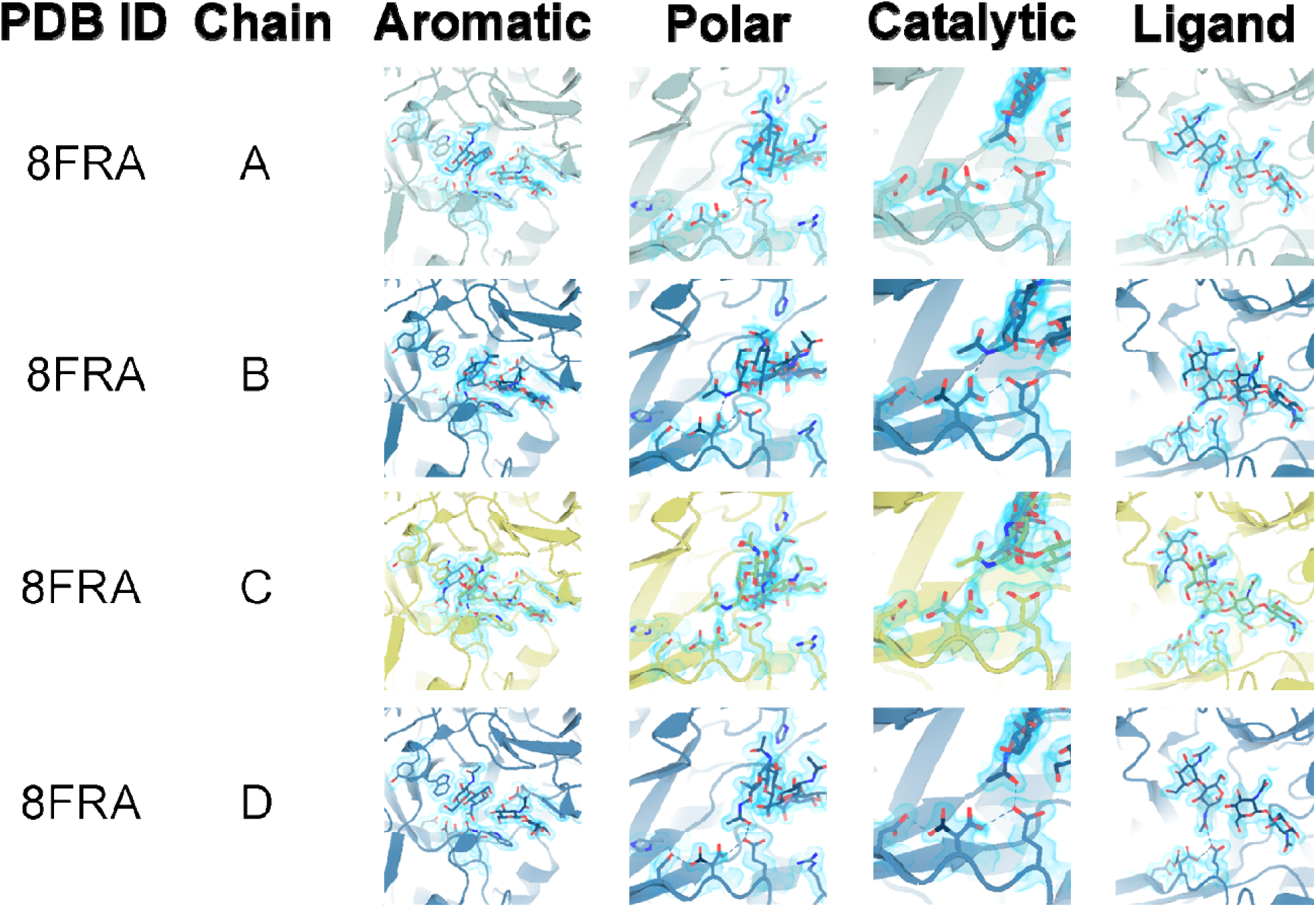
Overview of key residues for mAMCase activity. **A)** Stick representation of ligand and aromatic residues Trp31, Tyr34, Trp99, and Trp218 in the active site with 2mFo-DFc map shown as a 1.2 Å contour (blue). **B)** Stick representation of ligand and polar residues Arg145, His208, Asp213, and His269 in the active site with 2mFo-DFc map shown as a 1.2 Å contour (blue). **C, D)** Stick representation of ligand and catalytic residues Asp136, Asp138, and Glu140 in the active site with 2mFo-DFc map shown as a 1.2 Å contour (blue).

**Supplemental Figure 6.**
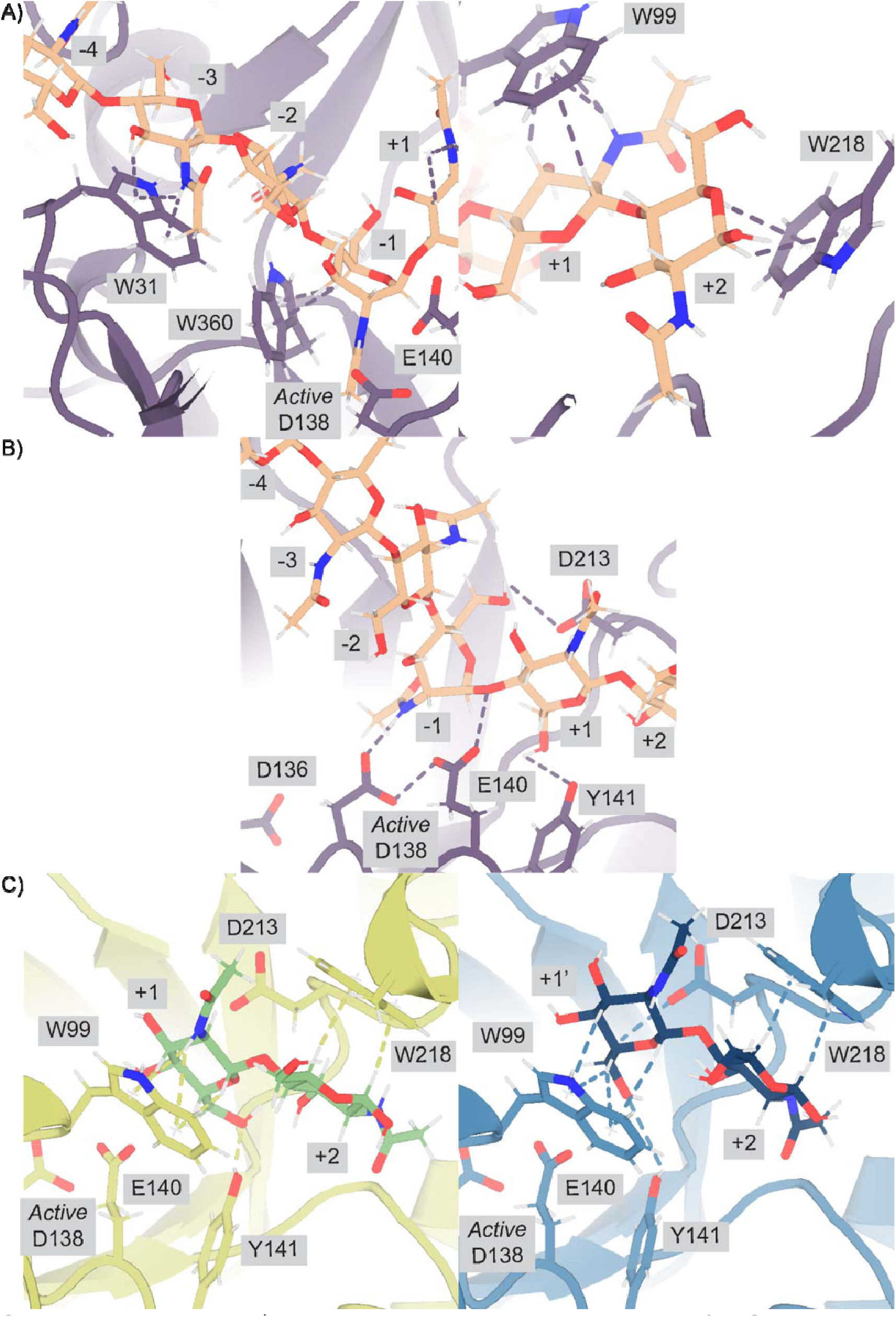
Protein-ligand interactions between mAMCase and chitin. **A)** PDB ID: 8GCA, chain A with GlcNAc_6_ modeled for viewing simplicity. Stick representation highlighting the stabilizing H-π interactions between Trp31, Trp360, and Trp218 and the -3, -1, +1, and +2 sugars, respectively. **B)** PDB ID: 8GCA, chain A with GlcNAc_6_ modeled for viewing simplicity. Stick representation highlighting the stabilizing hydrogen bond interactions between the -1 sugar and Asp138 (2.6 Å) and Asp213 (3.4 Å), and between the +1 sugar and Tyr141 (3.0 Å). Glu140 is 2.8 Å from the glycosidic oxygen bridging the -1 and +1 sugars. **C)** PDB ID: 8FRA, chains C (left) and D (right). Stick representation highlighting the stabilizing hydrogen bond interactions that we argue stabilize the +1 sugar (left; chain A) and the +1’ sugar-binding subsite (right; chain B).

**Supplemental Figure 7.**
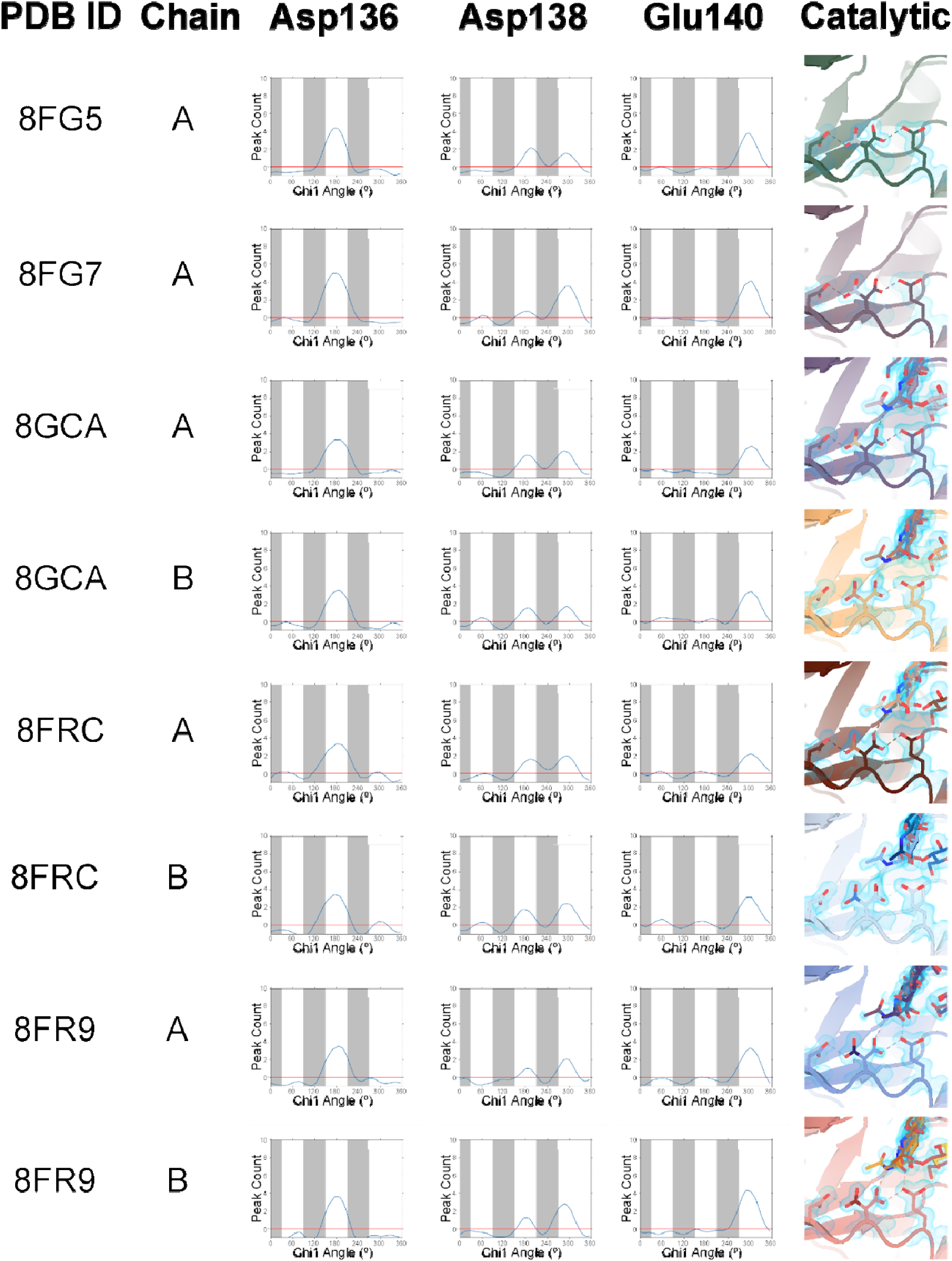
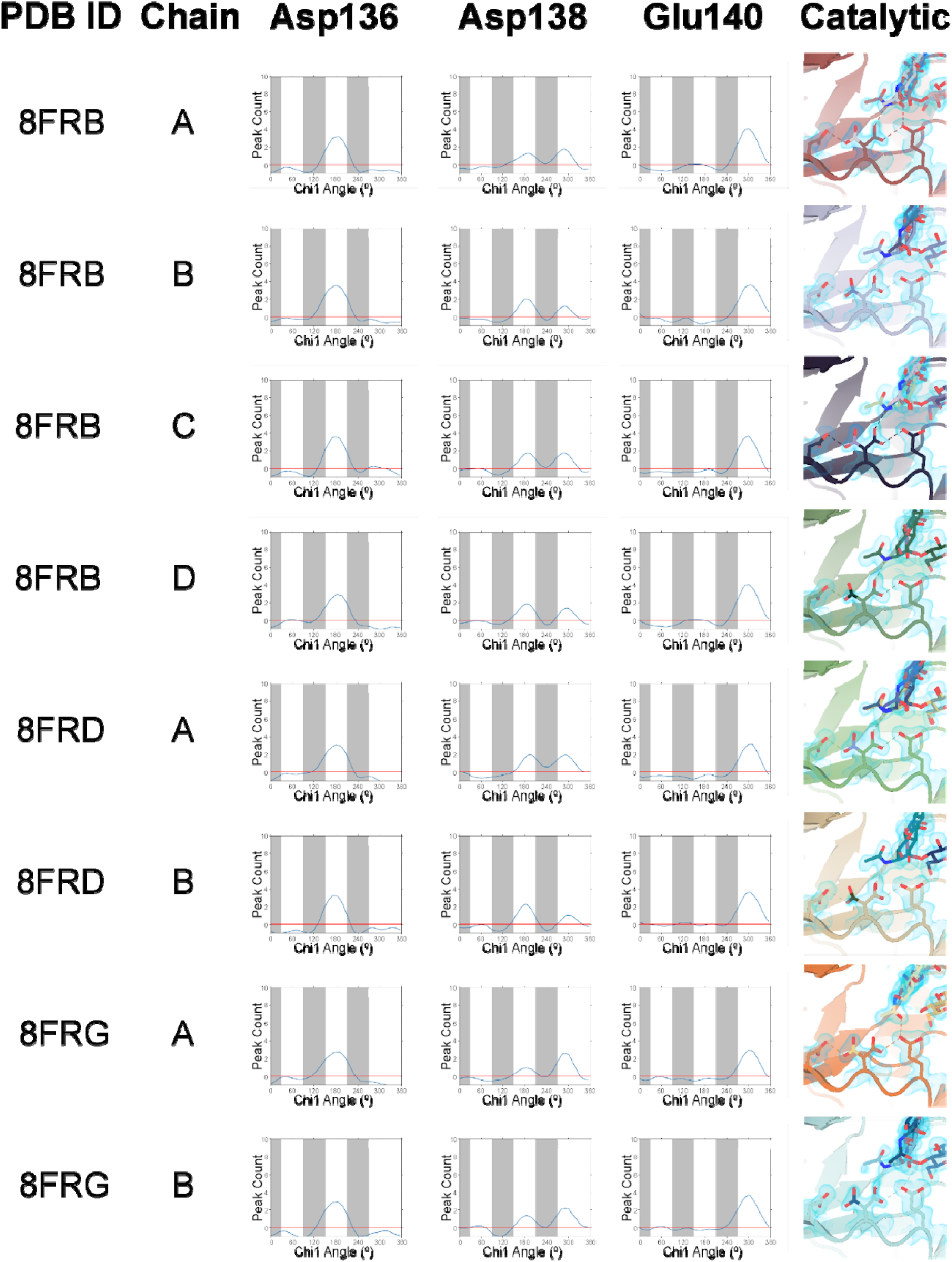
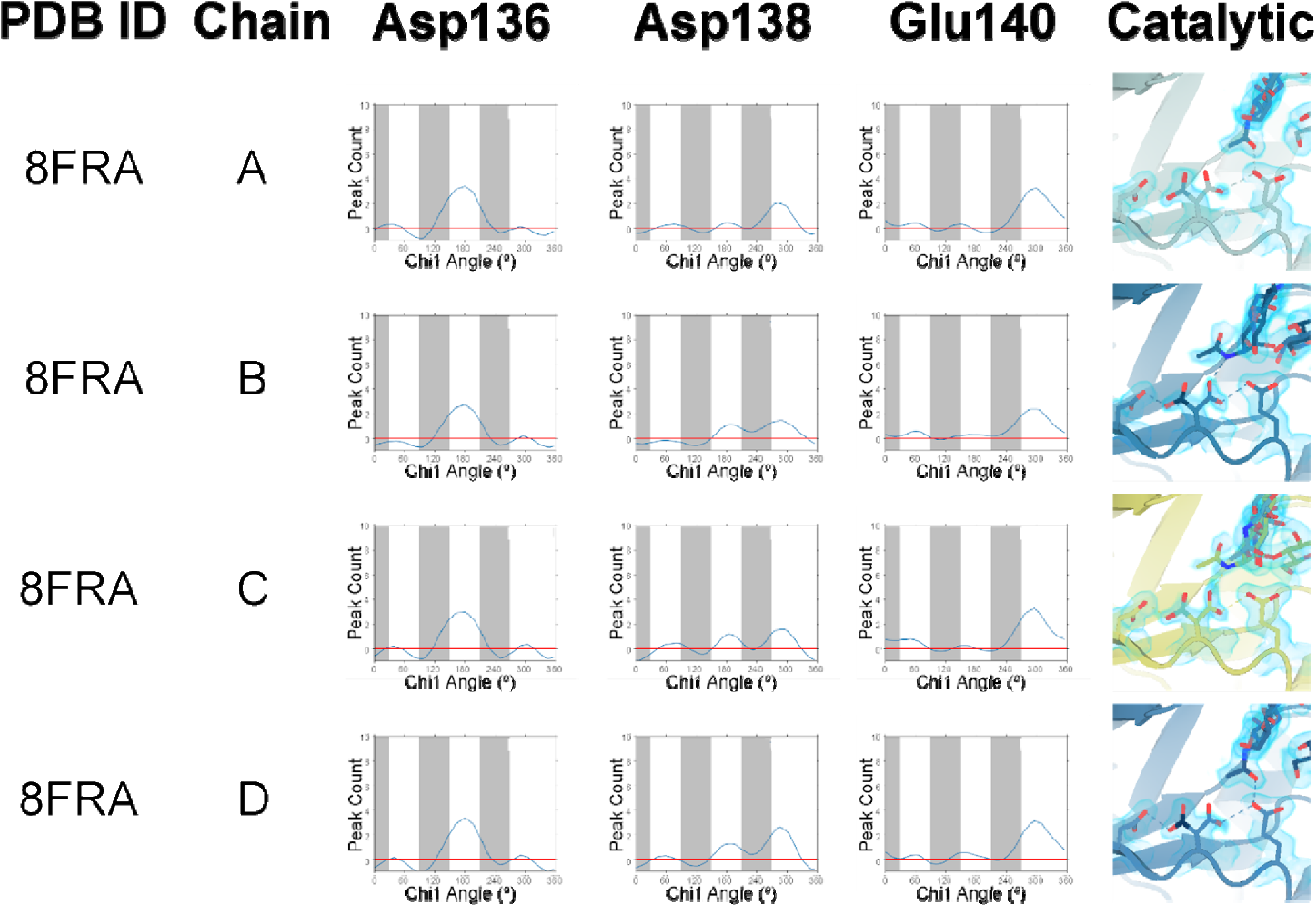
Ringer analysis of catalytic triad confirms alternative Asp138 conformations. **A)** Ringer analysis to detect alternative conformations in electron density maps. Ringer detected one peak for Asp136 at χ_1_ = 180° and Glu140 at χ_1_ = 300°, indicating only one conformation, whereas two peaks were detected for Asp138 at χ_1_ = 180° and χ_1_ = 300°, indicating two alternative conformations. **B)** Stick representation of Asp136, Asp138, and Glu140 with 2mFo-DFc map volume shown as a 1.2 Å contour (blue).

## General

We are grateful to Aashish Manglik and Mingliang Jin for providing ExpiCHO-S cells; to Liam McKay and Jose Luis Olmos, Jr. for support with the X-ray crystallography facility at UCSF; to George Meigs for assistance with X-ray data collection at ALS 8.3.1.; to Tzanko Doukov for assistance with X-ray data collection at SSRL 12-2; to Eric Greene, Duncan Muir, Stephanie Wankowicz, and Benjamin Barad for helpful discussions and critical feedback. Structural biology applications used at UCSF were compiled and configured by SBGrid.^55^

