## Supplementary figures and images for "Structural characterization of ligand binding and pH-specific enzymatic activity of mouse Acidic Mammalian Chitinase"

### SFigure 3.6B

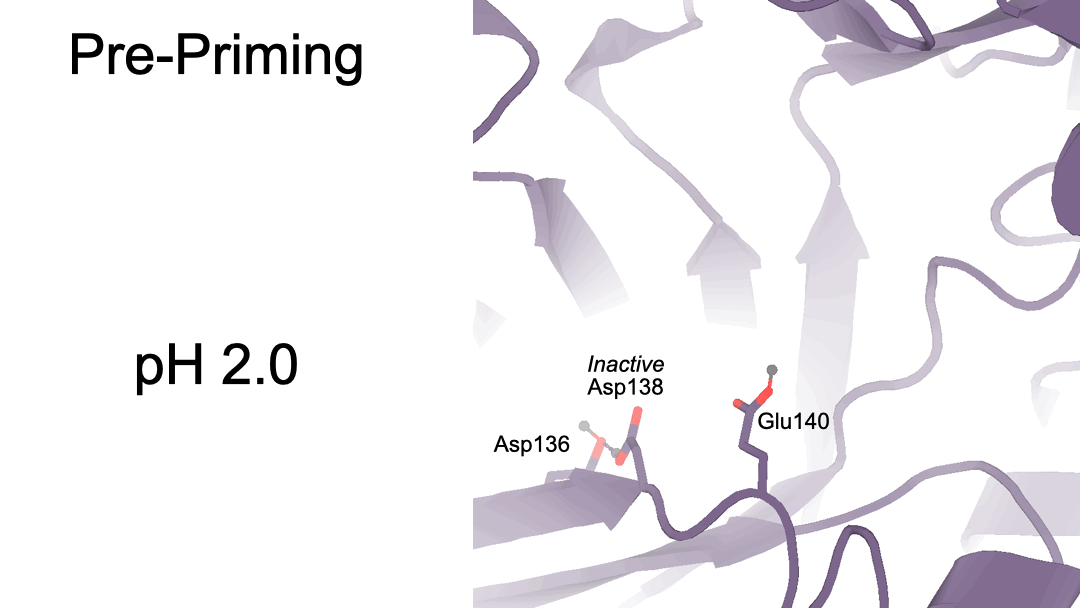

### SFigure 3.6C

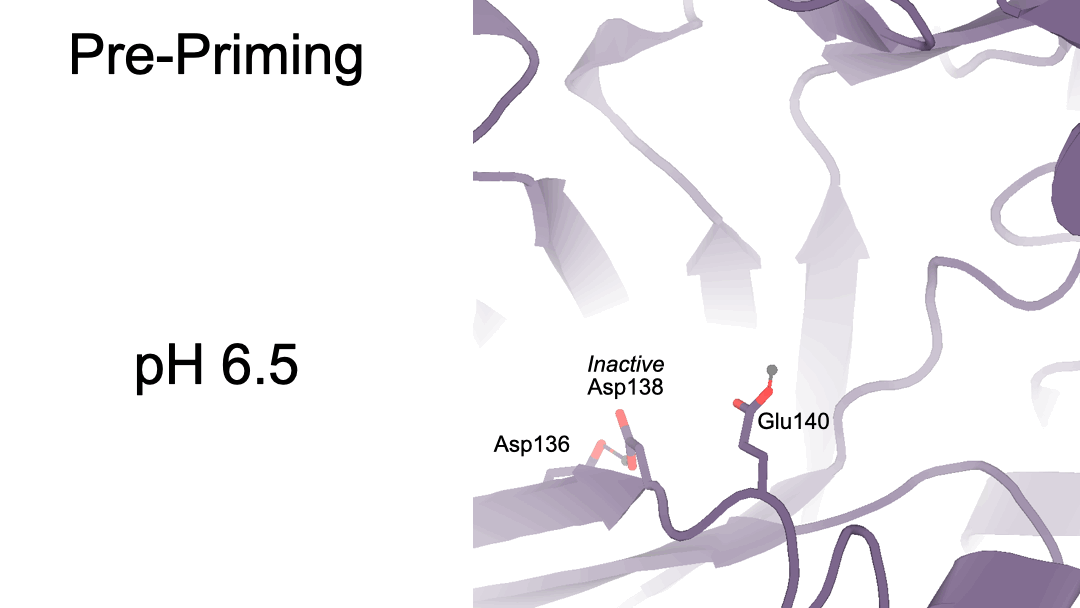
